# Missense mutations in CMS22 patients reveal that PREPL has both enzymatic and non-enzymatic functions

**DOI:** 10.1101/2023.12.18.572145

**Authors:** Yenthe Monnens, Anastasia Theodoropoulou, Karen Rosier, Kritika Bhalla, Alexia Mahy, Roeland Vanhoutte, Sandra Meulemans, Edoardo Cavani, Aleksandar Antanasijevic, Irma Lemmens, Jennifer A. Lee, Catherin J. Spellicy, Richard J. Schroer, Richardo A. Maselli, Chamindra G. Laverty, Patrizia Agostinis, David J. Pagliarini, Steven Verhelst, Maria J. Marcaida, Anne Rochtus, Matteo Dal Peraro, John W.M. Creemers

**Author notes:** Corresponding authors: John W.M. Creemers, Laboratory for Biochemical Neuroendocrinology, Department of Human Genetics, KU Leuven, ON1 Herestraat 49 – box 607, 3000 Leuven, Belgium. These authors contributed equally (Yenthe Monnens, Anastasia Theodoropoulou, Karen Rosier).

## Abstract

Congenital myasthenic syndrome-22 (CMS22) is a rare genetic disorder caused by mutations in the prolyl endopeptidase-like (*PREPL*) gene. Since previous reports only described patients with deletions and nonsense mutations in *PREPL*, nothing is known about the effect of missense mutations in the pathology of CMS22. In this study, we have characterized missense mutations in *PREPL* in three CMS22 patients, all with hallmark phenotypes. Biochemical evaluation revealed that these missense mutations do not impair hydrolase activity, thereby challenging the conventional diagnostic criteria. Structural analysis shows that the mutations affect regions most likely involved in intra-protein or protein-protein interactions. Indeed, binding to a selected group of known interactors was differentially reduced for the three mutants. The importance of non-hydrolytic functions of PREPL was investigated in catalytically inactive PREPL p.Ser559Ala cell lines which showed that hydrolytic activity of PREPL is needed for normal mitochondrial function but not for regulating AP1-mediated transport in the trans-Golgi network. In conclusion, these studies show that CMS22 can be caused not only by deletion and truncation of PREPL but also by missense mutations that do not necessarily result in a loss of hydrolytic activity of PREPL.

**Graphical abstract:** 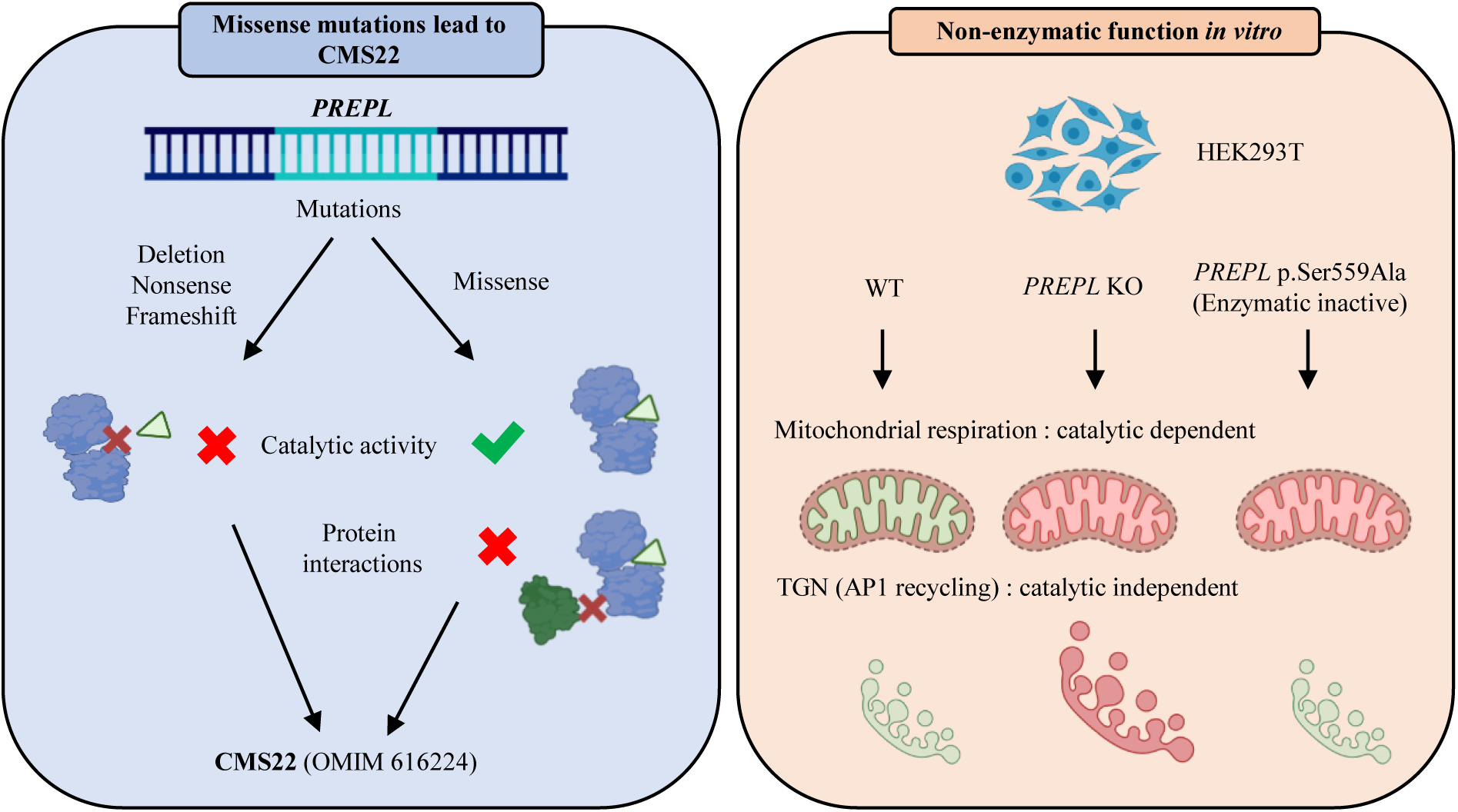

## Introduction

Congenital myasthenic syndrome-22 (CMS22, OMIM 616224) is a rare recessive disorder caused by mutations in the prolyl endopeptidase-like (*PREPL*) gene (1–6). Patients experience severe neonatal hypotonia, eyelid ptosis, feeding problems and growth hormone deficiency. During childhood, the hypotonia spontaneously improves and patients develop hyperphagia and rapid weight gain (1,2,4,6–13). Approximately half of the patients experience learning difficulties and have an average IQ of 70 (3). Previous reports have mostly documented CMS22 patient harboring partial or complete loss of *PREPL* alone or together with flanking genes (1,2,4,6–13). In addition, a small number of point mutations in the *PREPL* gene have been identified, leading to a frameshift or premature stop codon (1–6). Recently, a patient with a missense mutation in *PREPL* was reported but not functionally characterized(14).

The PREPL protein consists of two structural domains: an amino-terminal 7-bladed β-propeller domain and a carboxy-terminal α/β-hydrolase fold catalytic domain, connected by a hinge loop (7,15). The active site is comprised of the triad residues Ser559, Asp645 and His690. Two main isoforms of PREPL exist, a short form, PREPL_S_ (683aa), and a long form, PREPL_L_ (727aa). PREPL_S_ is localized in the cytosol while PREPL_L_ contains an N-terminal mitochondrial targeting sequence that translocates PREPL into the mitochondria (15–17). Experimental evidence confirms the catalytic activity of PREPL as it reacts with the activity-based probe FP-biotin, derived from the serine hydrolase inhibitor diisopropyl fluorophosphates, and can be inhibited by serine hydrolase inhibitors including PMSF and AEBSF (2,6,13). However, in contrast to its homologues PREP and OpdB, PREPL does not appear to possess protease activity despite the structural similarities (9,10). Recently, PREPL was found to hydrolyze chromogenic ester and thioester substrates in vitro and its activity was inhibited by Palmostatin M, a known inhibitor for acyl protein thioesterases 1 and 2 (APT1 and APT2) (15). Therefore, a depalmitoylating function has been proposed, as an increase in protein palmitoylation is evident in PREPL-deficient cells (18). Nevertheless, no physiological PREPL substrates have been identified to date.

In addition to its catalytic function, several lines of evidence support non-catalytic functions for PREPL. Firstly, yeast-two-hybrid experiments identified an interaction between PREPL and the µ1A subunit of the adaptor protein complex AP-1 (19). Notably, PREPL-deficient human skin fibroblasts showed prolonged membrane binding of AP-1 and a trans-Golgi network (TGN) expansion. This phenotype was rescued with a catalytically inactive PREPL p.Ser559Ala variant, suggesting a catalytic independent mechanism. Moreover, PREPL interacts with α-synuclein, thereby influencing aggregation, without α-synuclein cleavage (20). For PREP, a homologue of PREPL, substrate binding in the catalytic site is thought to result in an allosteric change that alters protein-protein interactions (21). It was further hypothesized that these interactions are physiologically most relevant, and that substrate binding and cleavage has only a modulatory role to the function of PREP. A similar mechanism of action could be valid for PREPL as well.

In this paper, we describe two new patients and one published but not functionally characterized CMS22 patient that harbor missense mutations in PREPL. We have studied the effect of the mutations on catalytic activity, protein stability, and protein-protein interactions. Experimental data demonstrate that compromising either the catalytic or non-catalytic functions of PREPL can result in CMS22. Using a HEK293T cell line harboring a p.Ser559Ala mutation in PREPL we investigated the need for catalytic activity of PREPL to maintain normal TGN morphology and mitochondrial respiration. Finally, we propose a new model for the physiological function of PREPL that integrates both its catalytic and non-catalytic functions.

## Materials and methods

### Bacterial protein expression and purification

The human PREPLs (Uniprot sequence Q4J6C6, residues 90-727) expression plasmid carries the gene for the PREPLs protein isoform, with an N-terminal 8xHistidine tag and an MBP (Maltose-binding protein) tag, followed by a TEV (Tobacco Etch Virus) cleavage site. PREPLs was cloned into the bacterial expression vector pVP68K and mutations were introduced using the QuikChange II site directed mutagenesis kit (Stratagene) and confirmed by direct Sanger sequencing. The plasmids were transformed into Rosetta(DE3)pLysS cells (Sigma-Aldrich). Protein expression was induced with isopropyl b-D-1-thiogalactopyranoside (IPTG; 0.4 mM) when cells reached an optical density (OD at 600 nm) of 0.9, with a subsequent growth over night at 20°C. Cell pellets were resuspended in lysis buffer (500mM NaCl, 50 mM Hepes pH 7.4, 1 mM TCEP, 5% glycerol) and were lysed by sonication. After centrifugation at 20,000xg for 60 min at 4°C, the supernatant was filtered and applied to two HisTrap HP columns (GE Healthcare). After elution of recombinant PREPL with a continuous gradient over 40 column volumes of elution buffer (500 mM NaCl, 50 mM Hepes pH 7.4, 1 mM TCEP, 5% glycerol, 500 mM imidazole), pure fractions were buffer exchanged into lysis buffer to remove imidazole, using a HiPrep Desalting column (GE Healthcare). The N-terminal purification tags were cleaved by overnight incubation with TEV protease. Subsequently, the cleaved tags and TEV protease were removed with another His affinity chromatography and the flowthrough was collected, concentrated, and additionally purified by Size Exclusion Chromatography (HiLoad 16/600 Superdex 200 pg, Cytiva) in lysis buffer. Pooled fractions were concentrated to approximately 10 mg/ml using a MWCO concentrator (Millipore) with a cutoff of 30 kDa, and stored at −20°C.

### Cell culture, transient transfection and immunoprecipitation

Generation of FLAG-tagged PREPL constructs for overexpression in mammalian cells has been described before (11). HEK293T cells were grown in DMEM supplemented with 10% FCS at 37°C, 5% CO2 and transfected using the Xtreme Gene 9 transfection reagent (Merck) according to the manufacturer’s protocol. 24 hours post-transfection, cells were lysed by passaging through a 26-gauge needle. Cell debris was removed by centrifugation at 16000g for 10 minutes. FLAG-tagged proteins were immunopurified using preformed complexes of anti-FLAG M2 antibody (Sigma Aldrich) and protein G Sepharose (GE healthcare).

### Metabolic labeling

Metabolic labeling was performed as described before (22). Briefly, transiently transfected HEK293T cells were starved for 1h in methionine-free RPMI (Thermo Fisher Scientific) and subsequently pulsed for 30 minutes in starvation medium supplemented with 200µCi/mL EasyTagTM EXPRESS 35S protein labeling mix (Perkin Elmer). Cells were chased in RPMI supplemented with 0.4 mM methionine for 0, 6, 12 and 24 hours, and subsequently lysed in 50 mM Tris pH 7.4, 150 mM NaCl, 1% Triton, 1 mM EDTA and 1 x cOmpleteTM mini protease inhibitor cocktail (Roche). PREPL proteins were immunoprecipitated using anti-FLAG M2 antibody (Sigma Aldrich) and separated by SDS-PAGE. Radioactive signal was amplified with NAMP100V (Sigma-Aldrich) and the gel was dried using a gel dryer (Biorad). Signal was captured using a TyphoonTM FLA 9500 biomolecular imager.

### Circular Dichroism Spectroscopy

Circular Dichroism Spectra of the PREPL variants (at 6 μM in 100 mM NaCl, 20 mM NaH2PO4 pH 7.4 and 1 mM TCEP) were collected with a Chirascan V100 (AppliedPhotophysics), using a 1 mm path length cuvette, at 20°C, in the 208-260 nm range. Thermal denaturation experiments were performed by heating the sample from 4 to 95 °C and collecting CD data at 217 nm every 2°C, with a temperature ramp speed of 1°C/min and an equilibration time of 15 sec.

### Isothermal titration calorimetry

Isothermal titration calorimetry (ITC) experiments were performed on a MicroCal PEAQ-ITC (Malvern Panalytical). PREPL proteins used for ITC measurements were in a buffer containing 300 mM NaCl, 20 mM Hepes pH 7.4 and 1 mM TCEP. Inhibitor 8 (1-isobutyl-3-oxo-3,5,6,7-tetrahydro-2H-cyclopenta[c]pyridine-4 carbonitrile) was synthesized and delivered as lyophilized powder (Vitas M Chemical) and was dissolved in 100% DMSO at a 100 mM concentration. Prior to the measurement, PREPL proteins were extensively dialyzed against the aforementioned buffer, and the ligand was diluted in the same dialysis buffer to prevent buffer mismatch, to a final concentration of 1 mM at 1% DMSO. Experiments consisted of titrations of a single 0.4 µL injection followed by 19 injections of 2 μl of titrant (inhibitor 8) into the cell containing PREPL protein at a 50 μM concentration.

Experiments were performed at 25°C in triplicate (n = 3), with injection duration of 4 s, injection spacing of 150 seconds, stir speed of 750 RPM, and reference power of 7 μcal/s. Data were processed and plotted using MicroCal PEAQ-ITC Analysis Software and fitted to a one-site binding model.

### Cryo-EM data acquisition

PREPL Arg243Cys was buffer exchanged into EM buffer (150 mM NaCl, 20 mM Hepes pH 7.4, 1 mM TCEP) by gel filtration chromatography using the Superdex 200 Increase 10/300 GL (Cytiva) and concentrated to 13 mg/mL using a 30 kDa cutoff MWCO concentrator (Millipore). Cryo-EM grids were prepared with an EM GP2 Automatic Plunge Freezer (Leica Microsystems). UltrAuFoil R1.2/1.3 Holey Gold foil on Gold 300 mesh grids from Quantifoil were glow-discharged for 90 s at 15 mA using the GloQube Plus Glow Discharge System from Quorum. 4.0 µL of PREPL Arg243Cys at 13 mg/ml was applied to the glow-discharged grids, back-blotted for 4 s under blot force 10 at 95% humidity and 4 °C in the sample chamber and plunge-frozen in liquid ethane, cooled by liquid nitrogen. Grids were loaded in a TFS Glacios microscope (200 kV) and screened for particle presence and ice quality, and the best grids were transferred to TFS Titan Krios G4. Cryo-EM data was collected using TFS Titan Krios G4 transmission electron microscope (TEM), equipped with a Cold-FEG on a Falcon IV detector in electron-counting mode (EER). Falcon IV gain references were collected just before data collection.

Data was collected using TFS EPU software package with aberration-free image shift protocol (AFIS), recording 9 micrographs per hole. Movies were recorded at nominal magnification of 250,000x, corresponding to the 0.3084 Å pixel size at the specimen level, with defocus values ranging from −1.5 to −2.5 µm. Exposure time was adjusted to 1 second per movie resulting in a total dose of 60 e-/Å^2^. In total, 10,745 micrographs in EER format were collected.

### Cryo-EM data processing

During data collection, initial processing was performed CryoSPARC live v3.2.0 (23) for assessing data quality. The obtained ab-initio structures from on-the-fly processing were used for particle picking and template generation. Motion correction was performed on raw stacks without binning, using the cryoSPARC Patch motion correction implementation. A total of 1,059,238 particles were template-based automatically picked and particles were binned by a factor of 3 during extraction. Two rounds of 2D classification were performed to remove bad particle picks, resulting in a set of 251,714 particles. Selected particles resulting from the 2D classification were used for ab initio reconstruction and subsequent non-uniform refinement and local refinement. After refinement, 137,436 particles were reconstructed into a 3D map of 4.01 Å resolution. The reported resolution is based on the gold-standard Fourier shell correlation 0.143 criterion. Local-resolution variations were estimated using cryoSPARC.

### Model building and refinement

The PREPL Arg243Cys model was generated using the crystal structure of PREPL WT (PDB 7OBM) (15), by mutating the Arg243 to Cys in UCSF ChimeraX (24). The generated mutant model was fit into the cryo-EM map with UCSF ChimeraX and was manually adjusted using Coot (25). The final model was generated after iterative cycles of manual model building in Coot and relaxed refinement in Rosetta (26). The areas corresponding to the N terminal loop of PREPL from residues 95-111 were not built, since the electron density at this area was not well defined. The same applies for the loop areas corresponding to amino acids 134-139 and 687-693. Sidechains of surface residues distributed all over the structure were deleted when their position was not well defined by the electron density. The final model was validated using MolProbity (27) and EMRinger (28) (Table S1). Figures were prepared using UCSF Chimera, UCSF ChimeraX and PyMOL.

### Activity-based probes (ABP) assay

Immunoprecipitated PREPL and PREPL variants from transiently transfected HEK293T cells were incubated for 30 min at 37°C with 1.5 µM FP-biotin in 50 mM phosphate buffer (pH 8) supplemented with 10 mM EDTA and 2 mM DTT. The reaction was stopped by the addition of SDS-PAGE loading buffer and samples were separated by SDS-PAGE. Covalent FP-biotin binding, indicative for serine hydrolase activity, was detected using a Streptavidine-HRP antibody (Dako). Membranes were subsequently stripped and relabeled with an anti-FLAG M2 antibody (Sigma Aldrich F3165) to detect the presence of PREPL protein for normalization.

### Fluorescent substrate cleavage assay

Substrate cleavage assays were done using 6,8-Difluoro-4-Methylumbelliferyl octanoate (DIFMUO), an octanoate esterified to a DIFMU fluorophore (29) DIFMUO substrate cleavage was performed using bacterially expressed PREPL variants as described above. Cleavage assays were performed in triplicate and each reaction consists of 110ul of reaction mixture containing 0.4 µM of protein, 0.1 mM DIFMUO, 50 mM Tris-HCl (pH 7.4), 150 mM NaCl and 9% DMSO. Fluorescence was measured at 1 min intervals for 100 min using a FLUOstar Galaxy microplate reader (BMG Labtech) at excitation and emission wavelengths of 390 nm and 460 nm respectively.

### MAPPIT evaluation of protein-protein interactions

MAPPIT was performed as described before (15,30,31) In short, HEK293T cells were plated in a 96-well plate at 10,000 cells/well. Subsequently, cells were co-transfected with 250 ng bait, 500 ng prey and 50 ng pXP2d2-rPAP1-luciferase reporter plasmid using the X-tremeGENETM 9 DNA transfection reagent according to the manufacturer’s protocol. A pSEL (+2L) PREPL_S_ containing vector was used as bait and screened with a selection of previously identified interactors in a pMG1 vector. Cells were stimulated 24 h after transfection using NeoRecormon® (Roche) at 5 ng/mL in DMEM while non-stimulated cells received DMEM. Luciferase activity was measured 24 h after stimulation using a Victor X3 plate reader (Perkin Elmer). Interactions were normalised using two negative controls: irrelevant bait (pSEL (+2L)) and irrelevant prey (pMG1).

### CRISPR-Cas9 genome editing

HEK293T PREPL knockout and PREPL p.Ser559Ala mutant cells were generated using the CRISPR-Cas9 genome engineering system according to Ran et al. (32). Briefly, guide RNAs targeting PREPL were designed and cloned into a pSpCas9(BB)-2A-Puro vector (Addgene 48139). For the p.Ser559Ala cell line, an additional repair ssODN template was designed containing the desired mutation. Subsequently, cells were transfected and treated with 2 mg/mL of puromycin for three days to select transfected cells. Single cells were grown in order to guarantee clonality. *PREPL* KO and PREPL p.Ser559Ala were confirmed using SDS-PAGE and Sanger sequencing. Oligonucleotides are listed in Table S2.

### FP-TAMRA activity detection of PREPL

HEK 293T cells (wild type, PREPL KO or PREPL p.Ser559Ala) were pelleted using a benchtop centrifuge (6000g, 10 min, 4°C). Pelleted cells were lysed by agitation in 150 µL lysis buffer (20 mM Na2HPO4, 750 mM NaCl, 0.5% Triton X-100, pH 7.4) in an Eppendorf tube. The lysates were spun down using a benchtop centrifuge (13000g, 5 min, room temperature) to pellet debris. Total protein concentration of each lysate was determined via BCA assay. All lysates were brought to the same protein concentration (1 mg/mL) by dilution with storage buffer (20 mM Na_2_HPO_4_, 750 mM NaCl, pH 7.4). The same storage buffer was used to make 5 µg/mL dilutions of purified PREPL proteins (wild type and catalytic mutant). 60 µL of all samples (HEK lysates and purified PREPL) was transferred to fresh Eppendorf tubes and incubated with 0.6 µL of a 100 µM stock of FP-TAMRA in DMSO (final concentration: 1 µM) for 1 hour. Next, 20 µL of 4x sample buffer was added to all samples, after which they were boiled at 95°C for 2 minutes. The samples were resolved on a 12% acrylamide gel using SDS-PAGE, after which probe-labeled proteins were visualized by measuring in-gel fluorescence on a Typhoon FLA 9500 scanner with excitation at 532 nm and detection at 568 nm with the photomultiplier set at 800 V.

### Confocal microscopy

Wild type and *PREPL* KO HEK293T cells were grown on glass coverslips coated with Poly-D-Lysine (Merck A-003-E) and fixed with 4% formaldehyde. Cells were labeled with anti-TGN46 antibody (Abcam) and Alexa Fluor-conjugated secondary antibody (Invitrogen). Cells were mounted using ProLongTM Antifade Mountant + DAPI (Thermo Fisher Scientific). Confocal images were captured using the Nikon C2 confocal microscope, with 60X oil immersion optics. Images were collected using the NIS Elements software (Nikon). Quantification was performed using ImageJ software (33).

### Seahorse cell mito stress test

Mitochondrial bioenergetics were analyzed by measuring oxygen consumption rates (OCR) using the Agilent XF24 extracellular flux analyzer (Seahorse Bioscience, North Billerica, MA) according to the guidelines of the manufacturer. Briefly, HEK293T were seeded in XF24 cell culture plates at a density of 45,000 cells/well and 30,000 cells/well in DMEM containing 10% FCS. After 20-24 hours incubation, adherent cells were washed twice with pre-warmed XF base medium supplemented with 10 mM glucose, 2 mM glutamine and 1 mM sodium pyruvate; pH 7.4 and incubated for 1h at 37°C. The sensor cartridge was hydrated overnight at 37°C with XF Calibrant and calibrated using the XF24 analyzer. Oxygen consumption rates (OCRs) were measured under baseline conditions followed by the sequential addition of oligomycin, carbonyl cyanide trifluoromethoxyphenylhydrazone (FCCP) and antimycin A to injector ports A, B, and C of the cartridge, respectively. The final concentrations of the injections were 2 mM oligomycin, 0.5 mM FCCP, and 1 mM antimycin A for HEK293T and 10 mM oligomycin, 2.7 mM FCCP, and 10 mM antimycin A for skin fibroblast cells. OCRs were normalized to the protein concentrations determined by the Pierce BCA Protein assay kit (Thermo Fisher Scientific).

### Western blotting

The PierceTM BCA Protein Assay kit (Thermo Fisher Scientific) was used to quantify protein concentrations. After denaturation, proteins were separated using SDS-PAGE electrophoresis on a 10% Bis-Tris gel and transferred to a nitrocellulose membrane. Next, membranes were blocked with 0.1M Tris HCl pH 7.4, 0.15M NaCl with 0.5% blocking reagent (Roche) supplemented with 0.2% Triton X-100. Membranes were labelled with primary antibody for 1 hour at room temperature or overnight at 4°C followed by incubation with HRP-conjugated secondary antibodies (Dako). Membranes were developed using the Western Lightning ECL system (Perkin Elmer). Antibodies : PREPL (Santa Cruz), FLAG M2 (Sigma Aldrich)

### Data analysis

Illustrations were made using Biorender.com. Image and westernblot analysis was performed using ImageJ software (33) Statistical analyses were performed using Prism GraphPad Prism 8 for Windows, GraphPad Software, Boston, Massachusetts USA, www.graphpad.com. Normal distribution of data was assessed with the Shapiro-Wilk test. For normally distributed data, statistical analysis was performed using one-way ANOVA and post-hoc Dunnett’s test for multiple comparison. For non-normally distributed data, statistical analysis was performed using the Mann-whitney U test with Dunnett’s multiple comparison. Value and error bars are represented as mean ± SD. Significance levels are shown as * p ≤ 0.05 ** p ≤ 0.01, *** p ≤ 0.001 and **** p ≤ 0.0001.

## Results

### 1) Three missense mutations in *PREPL* identified in CMS22 patients

Here we present two novel CMS22 patients with monoallelic missense mutations in *PREPL* combined with a monoallelic frameshift and early stop codon. In addition, a recent publication described a new CMS22 patient with a homozygous missense mutation in *PREPL* (34). An overview of patient characteristics is shown in Table 1.

**Table 1.** Patient characteristics.

|  | Patient 1 | Patient 2 | Patient 3 |
| --- | --- | --- | --- |
| **Genotype** | c.727C>T p.Arg243Cys,<br>p.Leu597Hisfs*4 | c.1235T>G p.Ile412Arg,<br>c.1978_1981DelCTCA<br>p.Leu660Argfs*22 | c.1940G>A<br>p.Arg647Gln, |
| **General** |  |  |  |
| Sex | F | M | M |
| Gestational age | 41 6/7 w | 39 w | 34 w |
| Birth weight | 3810 g (+0.07 SD) | 2825 g (-1.4 SD) | 1620 g (-4.39 SD) |
| Birth length | 51.4 cm (-0.23 SD) | 47 cm (-1.6 SD) | 41 cm (-4.69 SD) |
| **Endocrine** |  |  |  |
| Growth delay (age) | Yes (3y) | Yes (7y11m) | N/A |
| GH deficiency (age) | Yes | No | N/A |
| **Neuromuscular** |  |  |  |
| Neonatal hypotonia | Yes | Yes | Yes |
| Fluctuating weakness | Yes | Yes | N/A |
| Neonatal respiratory support | No | No | Yes |
| Age at independent walking (months) | 18 m | 24 m | Not possible |
| Nasal voice/dysarthria | No | Yes | N/A |
| Facial weakness | No | Yes | N/A |
| Ptosis | Yes | Yes | N/A |
| Residual weakness after age 3 y | Hypotonia, muscle fatigue and eyelid ptosis | Only nasal speech | N/A |
| **Gastro-intestinal** |  |  |  |
| Feeding difficulties | Yes (syringe feeding up to 2 m) | Yes | N/A |
| Transient tube feeding | No | N/A | N/A |
| Excessive weight gain (age) | No | Yes (6y) | N/A |
| **Cognitive** |  |  |  |
| Cognitive/behavioral problems | No | Yes, mild | Yes, severe developmental delay |
| **Other** |  |  |  |
|  | Mild myopathic facies<br>Dilated central canal from T8-conus, 2mm in diameter |  | Dysmorphic features, hearing disturbance, supraaortic stenosis, tethered spinal cord, cryptorchidism, duplex kidney. |
| **ΔΔG DynaMut** | 0.176 kcal/mol | -1.064 kcal/mol | -0,313 kcal/mol |
| **PolyPhen-2** | Benign | Probably damaging | Probably damaging |
| **SIFT** | Tolerated | Tolerated | Tolerated |

Patient 1 is a girl born by vaginal delivery at 41 weeks and 6 days post-induction. The pregnancy was unremarkable, although the fetus had reduced movements. She presented with severe neonatal hypotonia and feeding difficulties. Nasogastric tube feeding was started at 6 weeks of age. She presented intermittent mild ptosis but otherwise had a normal ophthalmologic examination, and a high-arched palate and no other facial weakness. She was very hypotonic, more pronounced in the upper extremities. Muscle tone was better distally in her upper extremities and in her legs. Deep tendon reflexes (DTRs) were decreased but plantar responses were bilaterally in flexion. She achieved independent sitting at 8 months and independent walking at 18 months. Brain MRI was normal but spine MRI demonstrated a syrinx (2mm diameter) from T8 to conus without evidence of myelopathy. Laboratory analysis showed a mildly low circulating insulin-like growth factor-I of 16ng/ml (ref 17-185 ng/ml) but with a normal insulin-like growth factor binding protein 3 (IGFBP3) for age, in which the former normalized over time without treatment. Urinary amino acids did not show cystinuria. She is now 4 years old and doing well on pyridostigmine and albuterol oral syrup. Current exam shows mild myopathic facies, high-arched palate, normal extraocular movements and facial expression, nasal speech, normal DTR, normal run, walk and rise from floor but with fatigue after high levels of activity only. She has steadily gained weight from the 1 percentile at birth to 36 percentile for age at 4 years of age. She was never treated with growth hormone. Language and cognitive development are normal and she is behaviorally and socially age appropriate. Sequencing identified a paternally inherited c.727C>T variant resulting in p.Arg243Cys and a maternally inherited duplication resulting in a frameshift and early stop codon p.Leu597Hisfs*4 in *PREPL* (NM 006036.4).

Patient 2 is a boy born at 39 weeks by Cesarean section for failure to progress after an uncomplicated pregnancy. His birth weight was 2825 g (−1.4 SD), length 47 cm (−1.6 SD) and head circumference 34 cm (−0.5 SD). He had profound neonatal hypotonia, poor sucking and feeding difficulties which persisted throughout infancy and childhood. He had slow weight gain and was underweight for his length in infancy. Starting from the age of 6 he had rapid weight gain. He started sitting and crawling at 12 months of age and walking at 24 months. He received physiotherapy, occupational therapy and speech therapy. He seemed weaker and more tired later in the day. He also sometimes had upper eyelid ptosis later in the day. He was diagnosed with obstructive sleep apnea during a pediatric nocturnal polysomnogram at 21 months of age. He was on CPAP/BiPAP until 3 years of age. Laboratory tests repeatedly showed normal IGF-I and insulin-like growth factor binding protein-3 (IGFBP3). Urinary amino acids did not show cystinuria. Currently he is 7 years and 11 months old with weight 24.7 kg (42nd percentile), height 116 cm (2nd percentile) and head circumference 51.7 cm (32nd percentile). He has nasal speech, normal facial expression, movement and strength. There is no ptosis. He has normal DTRs but an awkward tandem gait. He is social and learns well at school; there are no behavioral problems. Sequencing identified a paternal c.1235T>G variant resulting in p.Ile412Arg and a maternal c.1978_1981delCTCA variant leading to p.Leu660Argfs*22 in *PREPL* ((NM 006036.4).

Patient 3 was recently described by Kim et al. (2020) and was included in our biochemical analysis to characterize the pathogenicity of the missense mutation on PREPL functioning. The male patient is the first child of nonconsanguineous Korean parents. He was born at 34 gestational weeks with a birth weight of 1620 g (−4.39 DS), a height of 41 cm (−4.69 SD), and a head circumference of 30.5 cm (SD, −3.12 SD). Cesarean section delivery was performed because of oligohydramnios. Immediately after birth, he was admitted to the neonatal intensive care unit due to prematurity and respiratory distress. On physical examination at birth, dysmorphic features were noted, including thick arched eyebrows, hypertelorism, a broad prominent nasal bridge, low set ears, a cleft and high arched palate, and a short webbed neck. The patient showed global muscle hypotonia with limited spontaneous movement and decreased deep tendon reflex. The patient also had multiple additional anomalies, including hearing disturbance, supravalvular aortic stenosis, a tethered spinal cord, cryptorchidism, and duplex kidney.

The brain magnetic resonance image (MRI) at 1 month of age showed no specific abnormality. Laboratory tests, including a metabolic profile, plasma amino acids, urine organic acids, plasma acylcarnitines, and thyroid function were unremarkable. The karyotyping, multiplex ligation-dependent probe amplification (MLPA) analysis for 26 microdeletion syndromes, and the diagnostic exome sequencing of 4813 OMIM genes were also normal. At 2 years old, the patient presented global developmental delay, barely sat with support, and only spoke a single word. Whole exome sequencing identified a homozygous c.1940G>A variant resulting in p.Arg647Gln in *PREPL* ((NM 006036.4).

### 2) Catalytic activity is conserved in CMS22 patient-derived PREPL variants

These CMS22 patients have at least one *PREPL* allele that contains a missense mutation (p.Arg243Cys, p.Ile412Arg, p.Arg647Gln), located in the catalytic domain (Ile412 and Arg647) or the propeller domain (Arg243) (Fig. 1A). In order to provide a rational explanation for why these variants result in CMS22, we first investigated if they impair the catalytic activity, since all patients identified so far have a complete loss of PREPL activity. The three CMS22 mutants, wildtype (WT) PREPL and the catalytically inactive PREPL p.Ser559Ala (Fig. 1A) were produced in transiently transfected HEK293T cells, and also recombinantly expressed in bacterial cells. Recombinant WT PREPL, p.Arg243Cys, p.Arg647Gln mutants and the catalytically inactive p.Ser559Ala variant were soluble, while p.Ile412Arg PREPL was aggregated in vitro and could not be purified from bacteria. Transient expression of p.Ile412Arg PREPL in HEK293T cells seemed normal.

**Figure 1:**
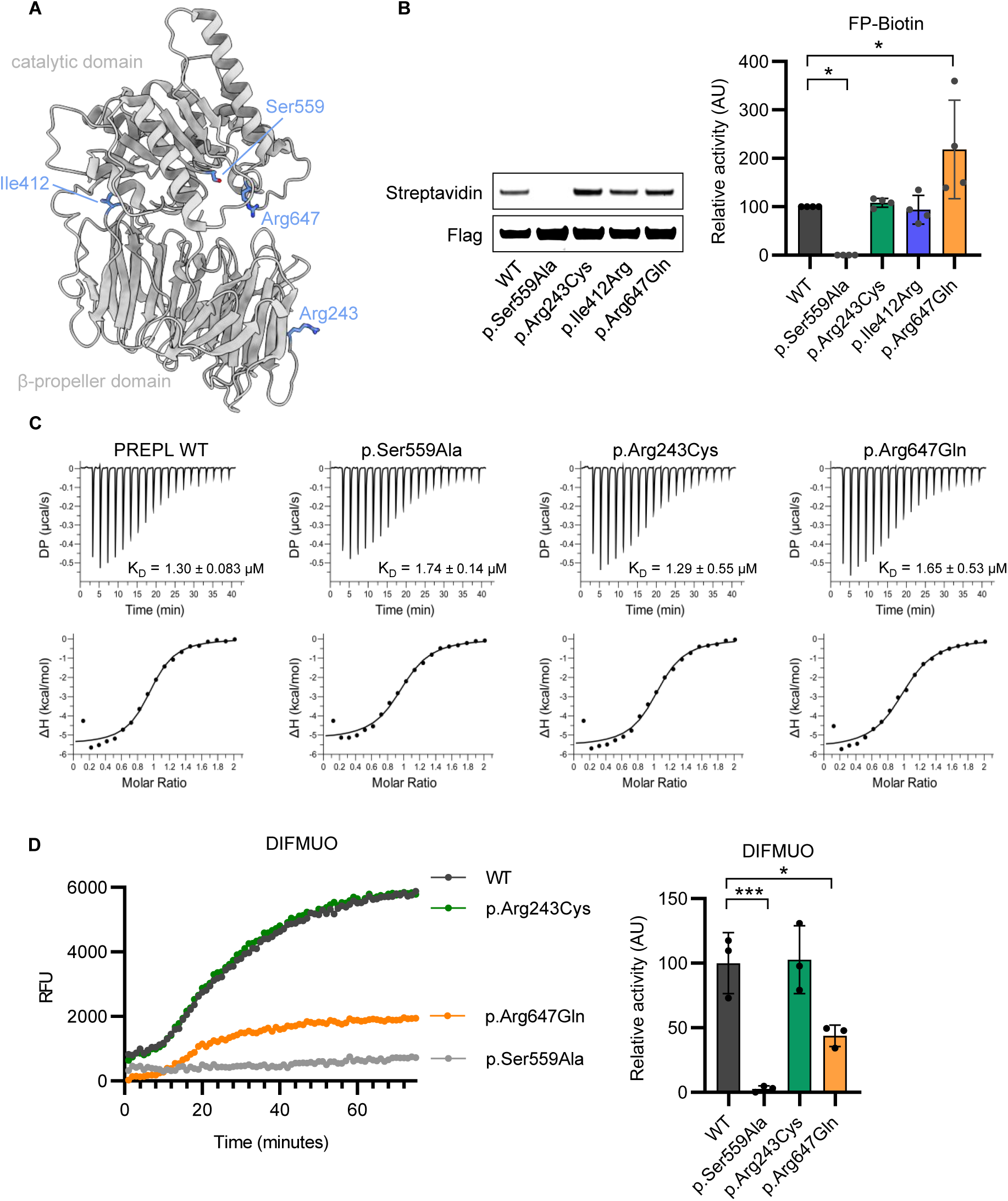
Catalytic activity of PREPL patient variants. **(A)** The three CMS22 mutants and the catalytically inactive PREPL p.Ser559Ala mapped on the PREPL structure. **(B)** Catalytic activity using FP-Biotin activity-based probe binding. **(C)** Isothermal titration calorimetry measurements of the binding of inhibitor 8 to WT, S559A, R243C and R647Q PREPL. Shown are single representative traces and each stated *K*_D_ value is the mean from n = 3 technical replicates. **(D)** Catalytic activity using DIFMUO substrate cleavage. Statistical analysis was performed using one-way ANOVA. Significance levels are shown as * p ≤ 0.05 ** p ≤ 0.01, *** p ≤ 0.001 and **** p ≤ 0.0001.

Using the serine hydrolase-specific activity-based probe (FP-biotin) we found that all three PREPL variants (p.Arg243Cys, p.Ile412Arg, p.Arg647Gln) were able to react with the probe, suggesting that the variants remain catalytically active (Fig. 1B). Furthermore, we observed that PREPL p.Arg647Gln (218.5 ± 101.6) had an increased affinity to the FP-biotin probe compared to WT (100 ± 0.0). The replacement of the positively charged Arg by the polar Gln is likely to facilitate the hydrophobic linker of FP-biotin, making this PREPL variant more reactive with the FP-biotin probe compared to WT. In addition, we found that the c.1978_1981/p.Leu660Argfs*22 deletion of patient 2 completely abolished the FP-biotin binding capacity of PREPL (Fig. S1).

To substantiate these results, we evaluated the binding dynamics of inhibitors to PREPL using Isothermal Titration Calorimetry (ITC). We tested the binding of inhibitor 8 (1-isobutyl-3-oxo-3,5,6,7-tetrahydro-2H-cyclopenta[c]pyridine-4-carbonitrile), a known inhibitor of PREPL (35) to the CMS22 PREPL variants. We found that WT PREPL, PREPL p.Arg243Cys, PREPL p.Arg647Gln and the catalytically inactive PREPL p.Ser559Ala bind inhibitor 8 with similar affinity, with an approximate value of 1 μM (Fig. 1C). This suggests that the binding between the protein and this inhibitor is dependent on the binding pocket and independent of the catalytic Ser559 residue. Moreover, binding remains unaffected by the mutations, which further supports the idea that the catalytic binding pocket is not affected by the mutations.

FP-biotin and inhibitor 8 binding only mimic the first steps of the PREPL enzymatic mechanism, i.e. formation of the Michaelis complex and – for FP-biotin – formation of a stable tetrahedral intermediate, but not the second aqueous hydrolysis step. Therefore, we also tested the ability of PREPL variants to cleave the fluorescent ester substrate 6,8-Difluoro-4-Methylumbelliferyl octanoate (DIFMUO) (29). This substrate could be cleaved by WT PREPL but not by PREPL p.Ser559Ala (Fig. S2). The substrate cleavage efficiency by WT PREPL was 50% compared to Acyl protein thioesterase 1(APT1) (Fig. 1D and S2). Moreover, we evaluated the cleavage of DIFMUO by CMS22 PREPL variants, where we saw that PREPL p.Arg243Cys (102.6 ± 26.2) had fully conserved catalytic activity, whilst PREPL p.Arg647Gln (43.8 ± 8.3) showed a twofold decrease in substrate cleavage activity compared to the WT (100.0 ± 23.6) (Fig. 1D). The proximity of the p.Arg647Gln mutation to the catalytic site could negatively impact the cleavage of this synthetic substrate.

The three point mutations described above, are the first missense mutations described in CMS22 patients, complementing the deletions, nonsense and frame-shift mutations. Since all three mutations have retained, at least partly, their hydrolytic activity we selected 18 PREPL missense mutations from the ClinVar database. The selection was based on their location in PREPL, thereby covering the whole protein, as well as positive selection if the variant was detected in a children’s hospital (36). From these, 8 reside in the β-propeller domain and 10 in the catalytic domain (Fig. S1A-C). Using the FP-biotin activity-based probe assay, we found that none of the mutations in the β-propeller domain had an effect on ABP binding. From the 10 mutations in the catalytic domain, 3 significantly decreased FP-biotin binding (p.Tyr479Cys, p.Arg510Gln, p.Ala560Pro) while 1 significantly increased FP-biotin binding (p.Glu605Lys). Ala560, but also Tyr479 are in close proximity with the catalytic Ser559, which explains the negative impact of the mutations on the ABP binding. Arg510 is localized at a distance from the catalytic triad residues, potentially explaining its less impactful decrease in ABP binding compared to the p.Tyr479Cys and p.Ala560Pro mutants. In conclusion, 15 out 18 tested variants in ClinVar have normal ABP binding. In the absence of case reports from the corresponding patients we cannot determine whether or not these are potential CMS22 patients, but it highlights the possibility of false-negatives if only ABP binding is used for differential diagnosis.

Moreover, since several premature stop codons have been found in CMS22 patients, we created 4 early stop codons at aa687, aa709, aa719 and aa724 to determine the minimal sequence for catalytic activity. Stop codons at aa719 and aa724 did not affect catalytic activity, while stop codons at aa687 and aa709 deactivated PREPL. It is worth noting that aa687 and aa709 reside in the loop preceding the last alpha helix and on the helix itself, respectively, within the PREPL structure. A stop codon mutation at these sites could disrupt the secondary structure of PREPL, rendering it catalytically inactive.

### 3) CMS22 PREPL mutants display similar stability to the WT protein

Since the catalytic activity of CMS22 mutants is conserved or slightly affected (in the case of p.Arg647Gln), we next sought to investigate the protein stability of these PREPL variants *in vitro*, to test if the CMS22 phenotype is a result of protein instability induced by the mutations.

Our results demonstrate that recombinant p.Arg243Cys and p.Arg647Gln PREPL mutants and the catalytically inactive p.Ser559Ala variant were soluble and displayed similar elution profiles to the WT protein by size exclusion chromatography, showing they are also monomeric (Fig. 2A). As already mentioned, p.Ile412Arg PREPL was aggregated *in vitro* and could not be purified. We next assessed the folding and thermal stability of PREPL mutants by circular dichroism (CD) spectroscopy. p.Arg243Cys, p.Arg647Gln and p.Ser559Ala PREPL share the same secondary structure as the WT protein (Fig. 2B). They all exhibited similar thermal stability (Fig. 2C), with the melting temperature (T_m_) of PREPL WT and the p.Ser559Ala, p.Arg647Gln mutants calculated to be 63.5 °C, while the melting temperature of p.Arg243Cys was 61.5 °C. Their unfolding process displayed a two-step pattern indicative of the presence of two domains as expected: the helical catalytic domain and the β-propeller comprised of beta sheets.

**Figure 2:**
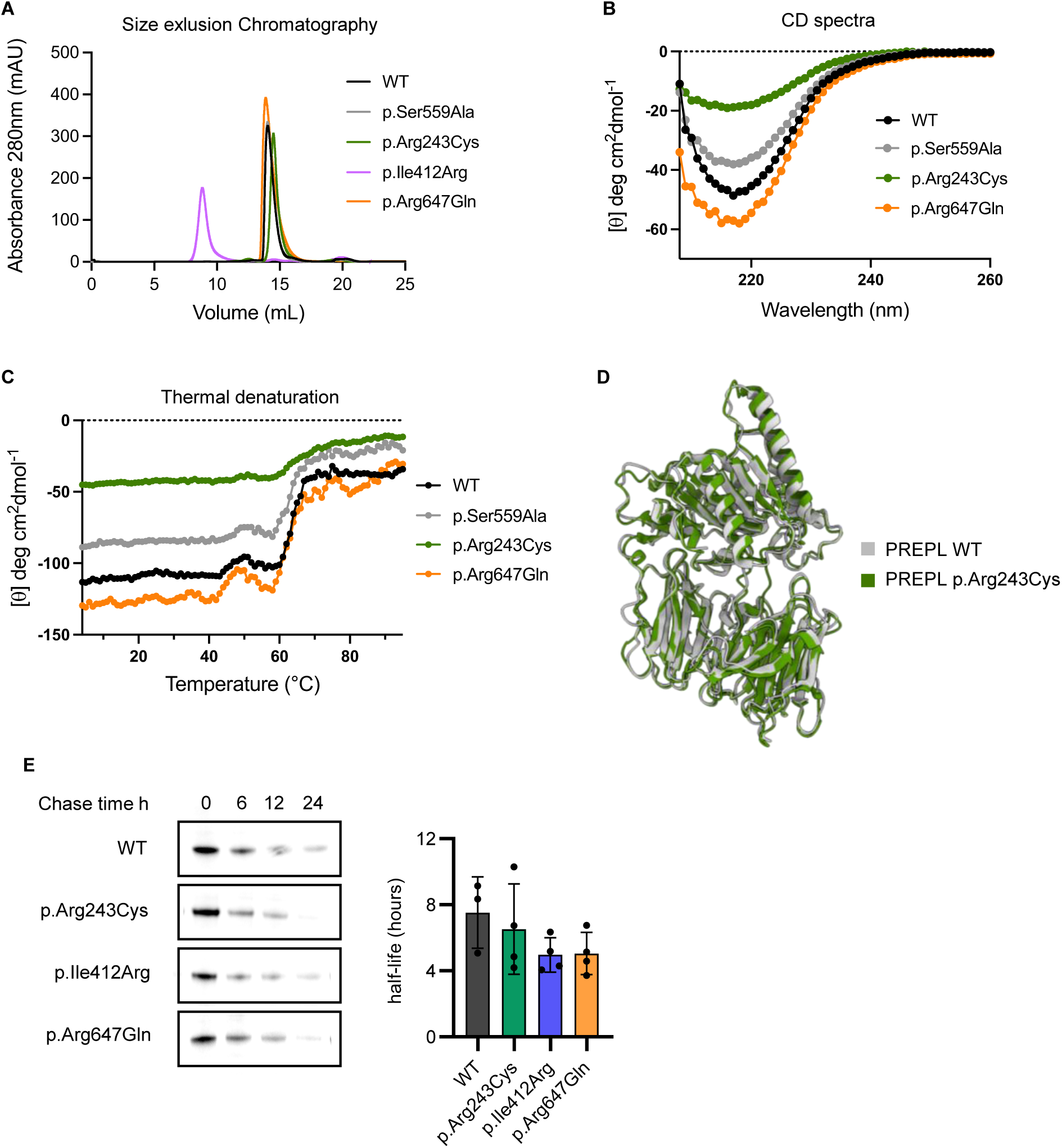
Protein stability of CMS22 mutants. **(A)** Recombinant p.Arg243Cys, p.Arg647Gln mutants and the catalytically inactive p.Ser559Ala PREPL variant are soluble and monomeric by size exclusion chromatogrpahy. Recombinant p.Ile412Arg PREPL mutant aggregates *in vitro*. **(B)** CD spectra of CMS22 mutants, which share the same secondary structure as the WT protein. **(C)** Thermal denaturation curves monitored by measuring the CD signal at 217 nm across a temperature range from 4 to 95°C. The melting temperature (T_m_) of PREPL WT and the p.Ser559Ala, p.Arg647Gln mutants is calculated to be 63.5°C, while the melting temperature of p.Arg243Cys is 61.5°C. The unfolding process of all PREPL variants displays a two-step pattern. **(D)** Structure of PREPL p.Arg243Cys mutant solved by cryo-EM in the open conformation, sharing the same fold as the WT protein. **(E)** Protein half life measured by pulse chase (value + blot).

To investigate the potential impact of the patient mutations on PREPL conformation we solved the structure of p.Arg243Cys mutant using cryo-electron microscopy (Fig.2D). This PREPL mutant was chosen due to its ability to be purified to higher concentrations compared to the other PREPL variants and its stable behaviour on the cryo-EM grids. The resulting map was resolved at a resolution of 4.01 Å (using the FSC 0.143 criterion) (Fig. S3). The novel structure of this CMS22 PREPL mutant shows that it is in an open conformation, highly-similar to the WT protein solved by crystallography (PDB ID: 7OBM) (Fig.2D), with a root mean square deviation (Ca) value of 0.980 Å. Therefore, this structural study shows that the Arg243Cys mutation did not affect the ternary structure of PREPL Next, we sought to evaluate protein half-life of the PREPL mutants *in vitro* in order to evaluate if the proteins are not subject to early degradation. Therefore, we performed 35S methionine labelling of HEK293T cells, transiently transfected with the PREPL variants to determine their half-lives. We established that WT PREPL has a half-life of 7.5 (± 2.2) hours (Fig. 2E). The half-lives of the patient PREPL variants were all slightly shorter, albeit non-significantly: PREPL p.Arg243Cys 6.5 (± 2.7) h, PREPL p.Ile412Arg 5.0 (± 1.0) h and PREPL p.Arg647Gln 5.1 (± 1.3) h. Overall, these results show that the mutations have limited or no effect on the stability of PREPL.

### 4) Patient mutations in PREPL disrupt protein-protein interactions

Recent findings suggest that protein-protein interactions are pivotal for the function of PREPL and that some of these interactions are independent of catalytic activity (15,19,20). Therefore, we evaluated if the CMS22 patient mutations have an effect on the interactome of PREPL. Using the mammalian protein-protein interaction trap (MAPPIT), protein-protein interaction scores were measured between WT and CMS22 PREPL mutants and 11 previously identified PREPL interactors (Fig. 3, Fig S4). These 11 interactors were selected from the previously identified list of 250 interactors, on the basis of a high interaction score, and divided between cell compartments and cell-biological functions (15). We found that patient-derived PREPL variants had significant changes in the interaction scores with the 11 interactors relative to WT PREPL. PREPL p.Arg243Cys had 7 decreased (NDUFV2, MRPL11, MRPS12, ARPC5L, DCTN5, DISC1, MDM1) and 1 increased interaction (UBL5), PREPL p.Ile412Arg had 8 decreased interactions (NDUFV2, MRPL11, MRPS12, STX5, ARPC5L, DCTN5, DISC1, MDM1) and PREPL p.Arg647Gln had 5 decreased (NDUFV2, MRPL11, STX5, ARPC5L, DCTN5) and 3 increased interactions (UBL5, ADM2, DISC1) (Fig. 3A, Fig. S4). These data indicate that the patient mutations alter the interactome of PREPL.

**Figure 3:**
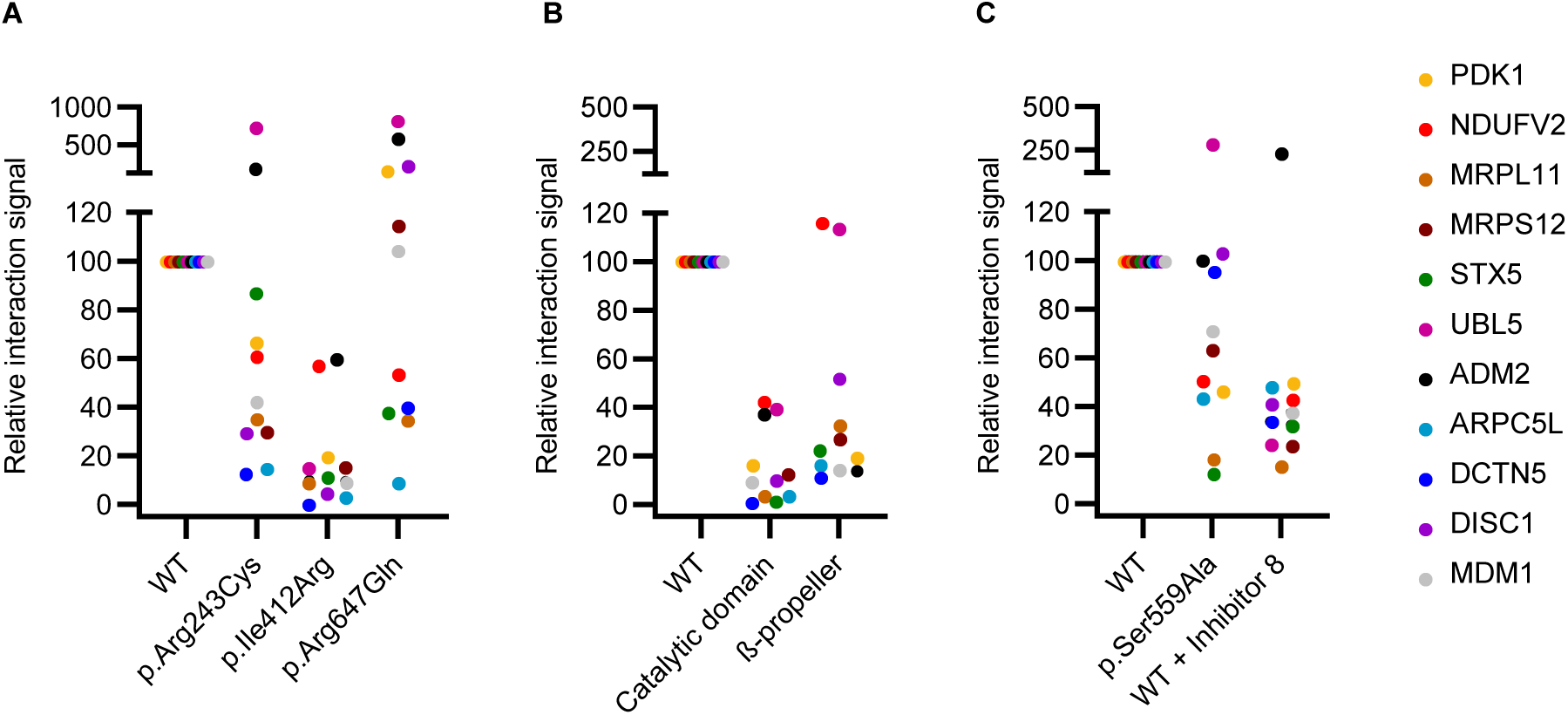
Overview of protein-protein interactions from 11 proteins with PREPL variants. Protein-protein interaction scores were determined using MAPPIT and relative interaction signal plotted for : A) WT PREPL, and PREPL variants p.Arg243Cys, p.Ile412Arg and p.Arg647Gln B) WT PREPL and the catalytic domain or β-propeller domain separately. C) WT PREPL and inactivated PREPL p.Ser559Ala or WT PREPL inactivated by addition of 50µM total concentration of compound 8.

To evaluate if protein-protein interactions of PREPL are domain specific we analysed the interactions formed by the catalytic domain and β-propeller domain independently (Fig. 3B, Fig. S4). We found that the individual domains were unable to maintain protein-protein interactions similar to WT. Only 2 interactors scored positively with the β-propeller alone (NDUFV2, UBL5). This indicates that both domains in the PREPL structure are needed for full protein-protein interactions, suggesting that interaction-sites span across both domains or are a result of a dynamic interplay between the two domains. Previously, Männistö et al. (2017) suggested that catalytic activity might have a modulatory role for allowing protein-protein interactions of PREP (21). Therefore, we evaluated if the catalytic activity contributes to protein-protein interactions by measuring interaction scores using a catalytically inactive PREPL p.Ser559Ala mutant and WT PREPL treated with the previously reported PREPL inhibitor 8 (Fig. 3C). Indeed in both cases, overall interactions were decreased with 6 interactors (NDUFV2, MRPL11, MRPS12, STX5, ARPC5L, MDM1) for the p.Ser559Ala mutant and 8 interactors (NDUFV2, MRPL11, MRPS12, STX5, ARPC5L, DCTN5, DISC1, MDM1) for the inhibitor 8-treated WT PREPL, respectively. This indicates that the catalytic activity is a contributing factor to protein-protein interactions of PREPL.

### 5) PREPL displays both catalytic and non-catalytic functions in HEK293T cells

Biochemical evaluation of the patient PREPL mutants shows that catalytic activity is conserved but protein-protein interactions are altered. Moreover, MAPPIT evaluation shows that both domains need to be present for protein-protein interactions and that catalytic activity might also play a contributing role in PREPL interactions. This can be as a modulatory function as proposed by Männistö et al. (2017) for PREP or else the measured interactor is a substrate on its own and binds in the substrate binding pocket (21). In order to evaluate the contribution of catalytic activity to the function of PREPL we used CRISPR-Cas9 to create 3 independent catalytically inactive PREPL p.Ser559Ala HEK293T cell lines, and one *PREPL* KO HEK293T cell line. After CRISPR-Cas9, Sanger sequencing revealed that all 3 PREPL p.Ser559Ala HEK293T cell lines have one mutant p.Ser559Ala PREPL allele combined with one early stop-codon allele. Using western blotting, we show in all 3 PREPL p.Ser559Ala/- cell lines, that PREPL was unable to bind to the fluorescent activity-based probe FP-TAMRA, thereby validating that PREPL p.Ser559Ala is catalytically inactive (Fig. 4A-B). These cell lines mimic the two patients with one null allele and one point mutation (Patient 1 & 2). PREPL activity was also recorded in WT, a PREPL+/- one allelic expression control cell line and recombinant expressed WT PREPL (Fig. 4B, red arrows).

**Figure 4:**
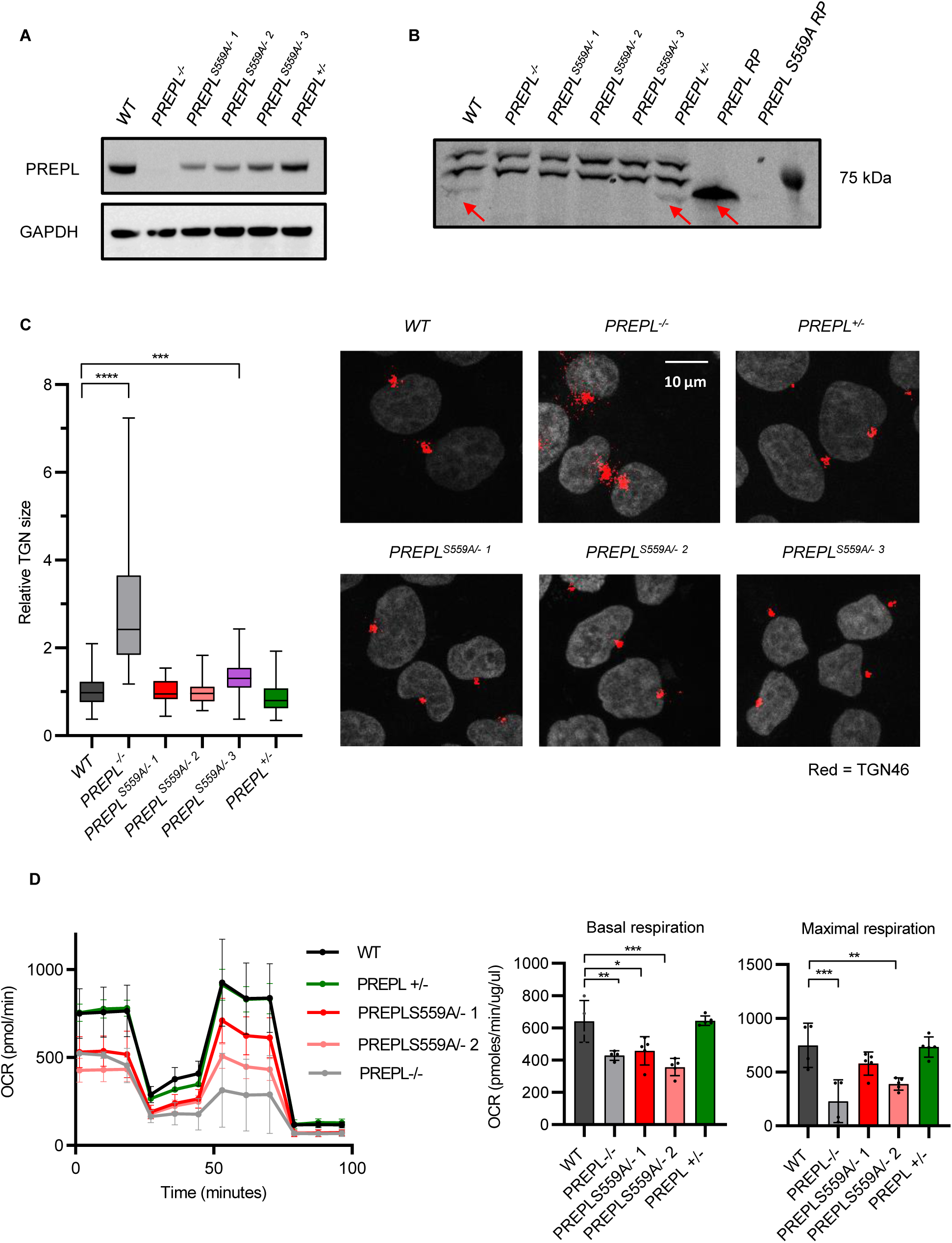
*PREPL*^S559A^ Mutant cells have a normal TGN morphology and display mitochondrial dysfunction. PREPL Knockout and p.Ser559Ala catalytic inactivated HEK293T cell lines were created using CRISPR-Cas9. A) PREPL expression levels in 20µM of HEK293T cell lysate. B) PREPL activity evaluated by FP-TAMRA. C) Evaluation of trans-Golgi network size by confocal microscopy. D) Evaluation of mitochondria function by seahorse cell mito stress test. Statistical analysis was performed using Mann-Whitney U test. Significance levels are shown as * p ≤ 0.05 ** p ≤ 0.01, *** p ≤ 0.001 and **** p ≤ 0.0001

To evaluate the contribution of catalytic activity to the function of PREPL, we subjected these PREPL p.Ser559Ala/- cells lines to two functional tests. First, Radhakrishnan et al. (2013) have demonstrated that PREPL interacts with the µ1A subunit of AP-1 and regulates AP-1 membrane binding (19). PREPL-deficient fibroblasts from hypotonia-cystinuria syndrome (HCS; MIM 606407) patients show an increased AP-1 membrane binding and present an increased TGN size. Increased AP-1 membrane binding could be rescued by overexpression of WT or p.Ser559Ala mutant PREPL, suggesting a non-enzymatic activity. We show that relative TGN size is significantly increased in PREPL -/- HEK293T cells (2.89 ± 1.45) compared to WT cells (1.00 ± 0.31) (Fig. 4C). Interestingly, two PREPL p.Ser559Ala/- HEK293T cell lines displayed a relative TGN size similar to WT: PREPL S559A/- 1 (1.00 ± 0.25) and PREPL S559A/- 2 (1.02 ± 0.33), whilst PREPL S559A/- 3 (1.34 ± 0.42) had a slightly increased relative TGN size. A normal relative TGN size was also recorded in the PREPL +/- one allelic expression control cell line (0.87 ± 0.34). Furthermore, addition of PREPL specific inhibitor 8 did not affect TGN morphology (Fig. S5). These data suggest that TGN morphology is regulated by PREPL in a catalytically independent manner and that monoallelic PREPL expression is sufficient to maintain normal TGN morphology.

Moreover, PREPL is involved in mitochondrial protein translation and mitochondrial respiration (15,17). Therefore, we evaluated mitochondrial respiration in our HEK293T cell lines using a Seahorse XFp extracellular flux analyzer (Fig. 4D). We found that basal mitochondrial respiration was significantly reduced in PREPL -/- HEK293T cells compared to WT whilst respiration remained normal in the PREPL +/- line. Moreover, reduced basal respiration was recorded in PREPL S559A/- 1 and PREPL S559A/- 2 HEK293T cells lines compared to WT cells. PREPL S559A/- 3 was not tested. Similarly, maximal respiration was significantly reduced in PREPL -/- and PREPL S559A/- 2 HEK293T cells compared to WT. These results suggest that catalytic activity is pivotal in the function of PREPL in the mitochondria.

## Discussion

In this study, we describe a new category of CMS22 patients, which should be taken into account in the differential diagnosis of congenital myasthenia. We studied three CMS22 patients that harbor compound heterozygous or homozygous missense mutations in *PREPL* that do not block the enzymatic activity of PREPL *in vitro*. These patients presented with the full CMS22 phenotype and share the hallmark phenotypes of neonatal hypotonia and growth delay. As CMS22 is a recessive disorder, these missense mutations must be detrimental to the function of PREPL in order to elicit the CMS22 phenotype. However, all three PREPL variants (p.Arg243Cys, p.Ile412Arg, p.Arg647Gln) react with the FP-biotin ABP. This shows that CMS22 diagnosis should not depend only on a negative FP-biotin test, thereby challenging the conventional diagnostic criteria (13). Moreover, PREPL p.Arg243Cys of patient 1 retained normal catalytic activity using a fluorescent substrate active *in vitro* while PREPL p.Arg647Gln only had a twofold reduced cleavage capacity.

Arg234 from the mutation of patient 1 p.Arg234Cys is positioned on PREPL in the loop connecting the β11 and β12 sheets of the third blade of the propeller and is not conserved across the homologous PREP, OpdB and AAP. It resides in an exposed region on the protein surface, where neither the original arginine nor the cysteine undergoes any interactions with neighboring residues (Fig. 5D). Despite the proximity of Cys227, the 9 Å distance between the two cysteine residues and the reducing environment of the cytoplasm makes the formation of a disulfide bond highly unlikely (Fig. S6A). Based on its location on PREPL structure, it is not surprising that the mutant retains catalytic activity but, instead, could very well affect protein-protein interactions mediated by the β-propeller.

**Figure 5:**
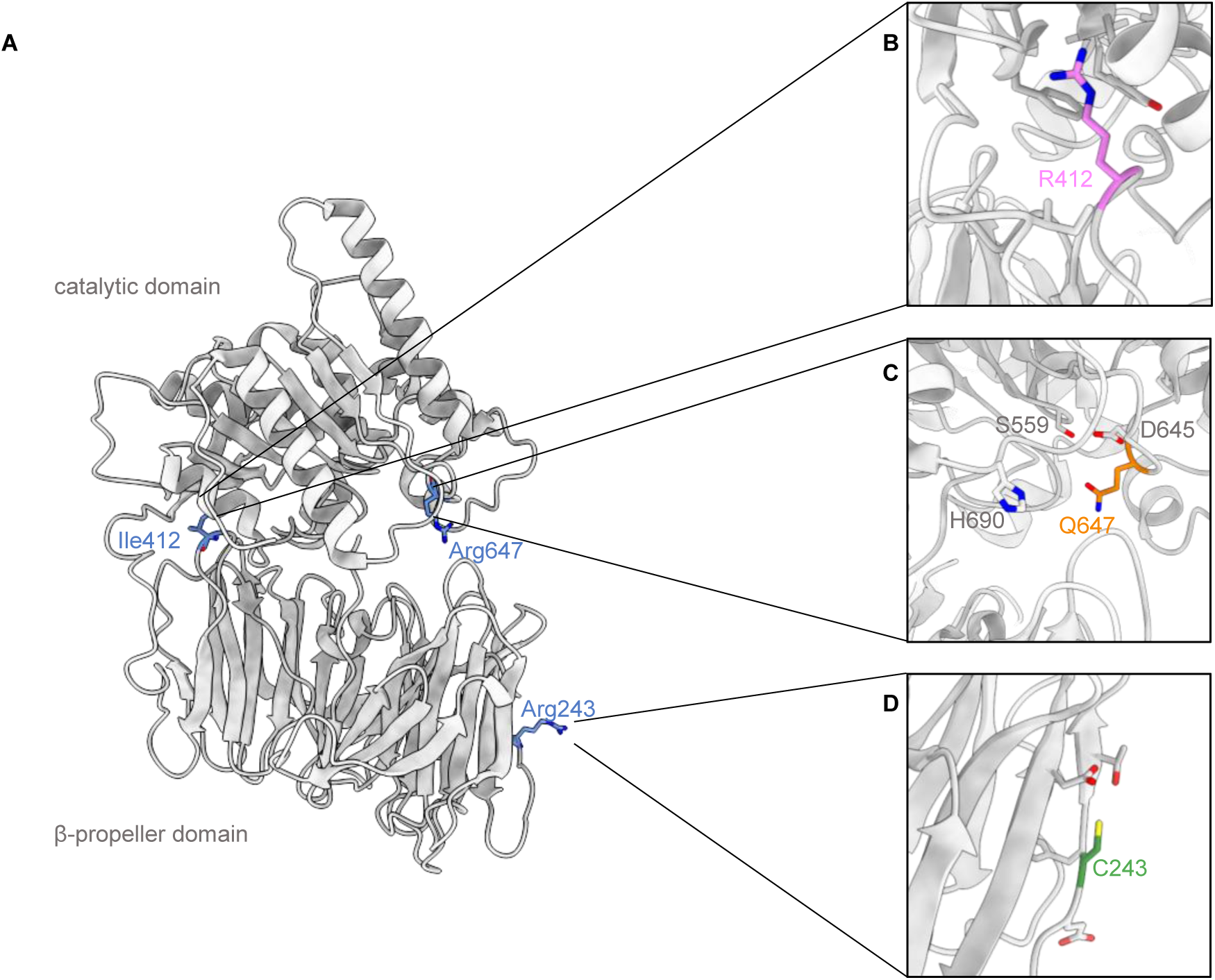
Location of CMS22 PREPL mutations on the PREPL structure and their effect on protein structure. **(A)** Location of the CMS22 patient mutations within the PREPL crystal structure. **(B)** Arg412 mutation on the β propeller in the vicinity of the hydrophobic residues Val498, Leu504, Ser441, Ile437, and Phe461. **(C)** Gln647 on the catalytic domain close to the catalytic triad residues Ser559, Asp645 and His690. **(D)** Cys243 and nearby PREPL residues at an exposed surface of the β-propeller domain.

p.Ile412Arg in patient 2, is located on the β-propeller loop that links the β 26 and β 27 sheets of the last blade, just before the hinge loop (residues 424-440) that connects the two PREPL domains (Fig. 5A) and does not display conservation within homologous protein sequences. Ile412 faces the catalytic domain of PREPL and resides in a hydrophobic region of the protein formed by Val498, Leu504, Ser441, Ile437, and Phe461 (Fig. S6B). The substitution of the hydrophobic isoleucine with the positively charged arginine could potentially disrupt this hydrophobic pocket within PREPL (Fig. 5B). This could consequently impact the interaction between the two domains, potentially leading to protein instability, causing aggregation *in vitro*.

Lastly, Arg647 from the mutation of patient 3 p.Arg647Gln, is a highly conserved residue across PREP, OpdB and AAP, and is positioned on a loop within the catalytic domain of PREPL, in close proximity to the catalytic triad residues Ser559, Asp645 and His690 at the interface between the two domains (Fig. 5C). It has previously been proposed to participate in a putative lipid-binding pocket in PREPL (15). In the WT PREPL structure, Arg647 is found in near proximity to Glu603 and Glu646 of the catalytic domain, implying a potential interaction between these residues (Fig. S6C). Additionally, Glu101, Glu106 and Glu107, located on the missing loop extending from the amino terminus, could potentially interact through the formation of salt bridges with Arg647. Hence, the introduction of the Arg647Gln mutation could disrupt these interactions, affecting the structure and dynamics of the protein.

Given the conservation of Arg647 among homologous proteins, we investigated the significance of this residue across these protein structures. In the closed conformation of OpdB (PDB 4BP9), Arg650 (Arg647 in PREPL) forms a salt bridge with the β-propeller residue Glu172 (Glu180 in PREPL based on structural superposition), stabilizing the closed active conformation (Fig. S6D). In the open conformation of OpdB (PDB 4BP8) Glu172 (Glu180 in PREPL) is within a 16 Å distance from Arg650, rendering their interaction highly unlikely. In this case, Arg650 interacts with Asp648 in OpdB. The same is observed in the closed conformation of PREP (PDB 3IUL) where Arg624 interacts with Asp622 of the catalytic domain (Fig. S6E) The multitude of interactions observed suggests the importance of Arg647 across PREPL homologues and provides an explanation as to why its mutation contributes to CMS22. We showed that all CMS22 PREPL variants have a normal thermostability and (near-)normal half-lives. Moreover, as suggested by the p.Arg243Cys cryoEM structure that we were able to solve, they could also share the same open conformation. Nevertheless, the interactome of the PREPL variants was severely affected.

The two patients examined in this study were compound heterozygous with one allele containing a frameshift mutation leading to a premature stop codon. In both cases, the carboxyterminal deletion (at Leu597 and Leu660) leads to a catalytically inactive allele since we demonstrated that truncation of PREPL at Glu687 already inactivates PREPL. These stop codon mutations disrupt the secondary structure of PREPL by cutting the last helix of the catalytic domain short, and lead to a catalytically inactive PREPL. The third patient was homozygous for p.Arg647Gln, a mutation that reduced the enzymatic activity and altered its specificity. How big the impact is on physiological substrates remains to be established.

In contrast to the limited effect on the catalytic activity of PREPL, its MAPPIT interactome was more dramatically altered. MAPPIT is a well-established mammalian two-hybrid interaction mapping technique capable of identifying unique interaction partners with a high signal-to-noise ratio due to effective controls (empty prey, empty bait) to account for non-specific signaling (30,31). Using 11 previously established mitochondrial and cytoplasmic MAPPIT interactors, we found that the patient missense mutations p.Arg243Cys and p.Ile412Arg significantly reduced the interactions with (almost) all interactors. Ipso facto, loss of interactions is the likely cause for loss of function of this allele in patient 1 (p.Arg234) and 2 (p.Ile412Arg). Remarkably, the Arg647Gln showed reduced interactions with about half the interactors, but increased interactions with the other half. The increased reactivity with FP-biotin but decreased cleavage of the DIFMUO ester substrate are consistent with a model in which this mutation results in reduced activity and altered specificity ultimately leading to a null allele. Further *in vitro* studies have highlighted the importance of both domains being together to facilitate protein-protein interactions and the necessity of functional catalytic activity, hinting at a collaborative function between catalytic activity and protein-protein interactions.

To shed more light on the collaborative function between catalytic activity and protein-protein interactions, we evaluated the effect of catalytic inactivation (p.Ser559Ala) on the cell-biological function of PREPL. Mitochondrial respiratory activity is dependent on catalytic activity of PREPL, suggesting that substrate cleavage is crucial for mitochondrial regulation. Since the missense mutations in patient 1 (p.Arg234) and 2 (p.Ile412Arg) are not altering the catalytic activity of PREPL active, we expect mitochondrial dysfunction due to the loss of protein interaction prior to substrate cleavage. Previously, Radhakrishnan et al. have shown that TGN morphology is regulated by PREPL independent of catalytic activity through protein-protein interaction with the µ1A subunit of AP1. We created a more robust and high throughput method to measure TGN size with increased sample size and used endogenously inactivated PREPL p.Ser559ala in order to validate that AP1 recycling does not require the catalytic activity of PREPL. Together, these functional assays highlight the importance of catalytically independent protein-protein interactions for the physiological function of PREPL.

Based on our results, we propose a new model for PREPL function (Fig. 6). In this model, PREPL first interacts with a target protein during which substrate specificity is determined. Possible alignment of a posttranslational modifications (PTM) like palmitoylation within the substrate binding pocket of PREPL can be the cause or consequence of the protein interaction. Subsequently, two different paths are possible: 1) PREPL cleaves a PTM from the target protein, thereby completing its physiological function and ending the protein-protein interaction. 2) PREPL continues the protein interaction and takes part in complex formation. This can either be reversible, requiring no need for catalytic activity, or the protein interaction can be terminated by cleavage of a PTM of the target protein. In this functional cycle, both a catalytic and non-catalytic mechanism of action of PREPL can coexist or collaborate to facilitate the physiological function of PREPL. Contrary to the previously suggested functional cycle, catalytic activity does not modulate protein-protein interactions but can be part of the interaction itself.

**Figure 6:**
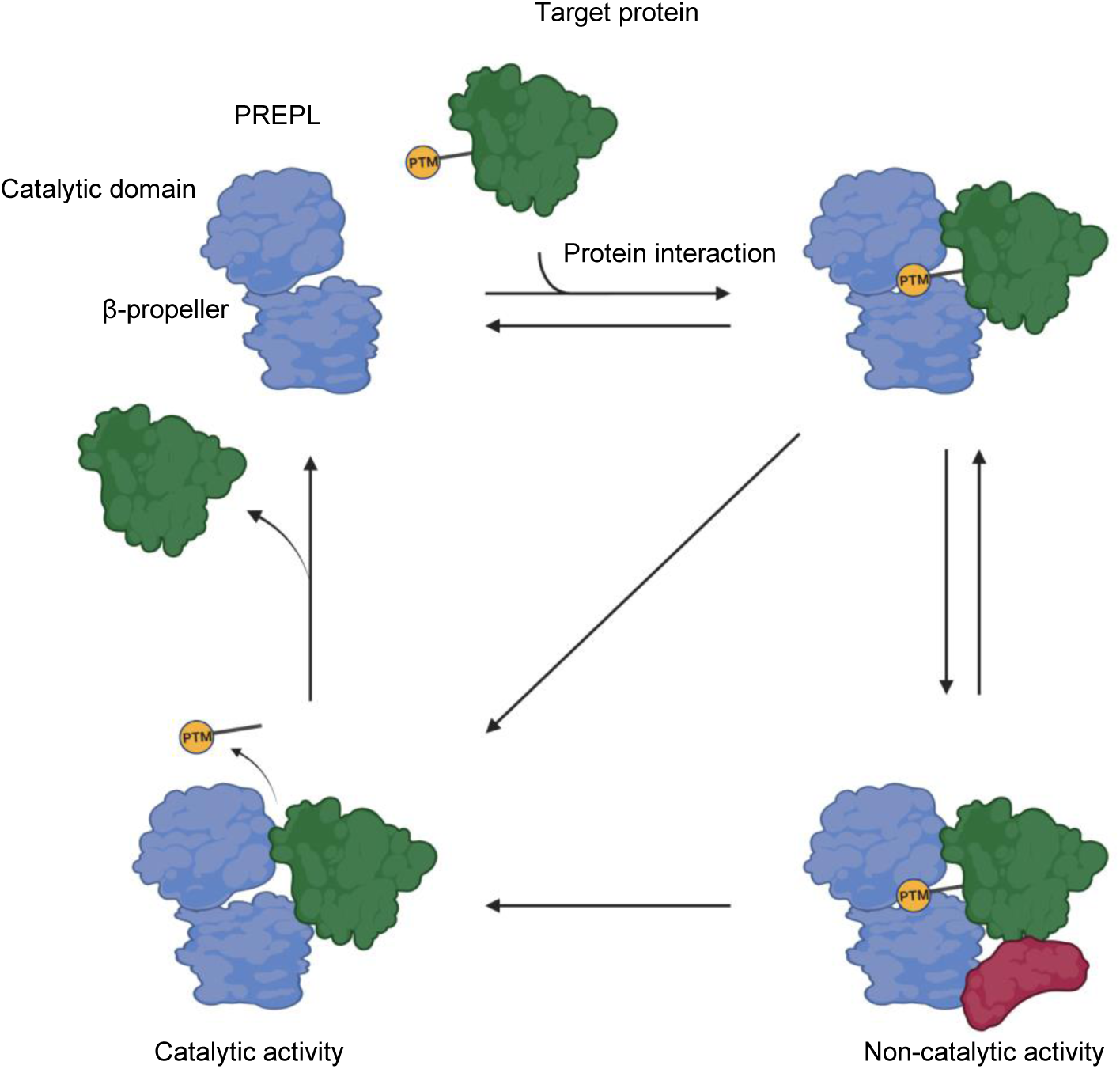
A proposed functional cycle for PREPL incorporating catalytic and non-catalytic activity. Based on our in vitro results, we propose a novel functional cycle for PREPL. In this cycle, PREPL interacts with a target protein via a protein-protein interaction. This interaction determines substrate specificity. Alignation of a post-translational modification inside the substrate binding pocket of PREPL can be part or consequence of the interaction. Non-catalytic activity can be mediated through reversible complex formation whilst catalytic activity consist of PTM cleavage form the target protein.

In conclusion, in this study we have presented a new category of CMS22 patients. This new category of patients still has PREPL hydrolytic activity *in vitro* but impaired protein-protein interaction, leading to impaired function and full-spectrum CMS22. Our previously developed functional blood assay (13) remains useful for identifying deleterious mutations in intronic or regulatory genomic regions, but exonic variants require further scrutiny. In addition to reactivity to an ABP or synthetic substrate, protein-protein interactions will have to be considered.

## Data availability

The reconstructed map of PREPL Arg243C has been deposited to the Electron Microscopy Data Bank (EMDB) with the accession number EMD-19117. The atomic model is deposited in the Protein Data Bank (PDB) with accession number PDB-8RFB. The raw cryo-EM images will be deposited in the EMPIAR database.

## Acknowledgements

We thank the VIB Bio-imaging core for use of confocal microscopy and the KU Leuven genomics core for plasmid DNA sequencing. We thank the KU Leuven C14/21/095 InterAction consortium for the purchase of Seahorse Analyzer used for mitochondrial studies and Ellen Vervoort for her support with the Seahorse XFp extracellular flux analyzer. We thank the EPFL Protein Production and Structure Core Facility for providing the equipment for the biophysical characterization of the proteins; Yoan Duhoo and the staff members of the Dubochet Center for Imaging in Lausanne, in particular Emiko Uchikawa and Sergey Nazarov, for their assistance with cryoEM sample preparation and data collection and analysis; Kelvin Lau and Luciano Abriata for helpful discussions and advice. This work was supported by FWO project #GOB9119N, FWO fellowships 1SE4424N and 12A2723N, the EPFL and the Swiss National Science Foundation (200021-157217 to MDP).

## Conflict of interest statement

The authors have declared that no conflict of interest exists.

---

**Table S1.**
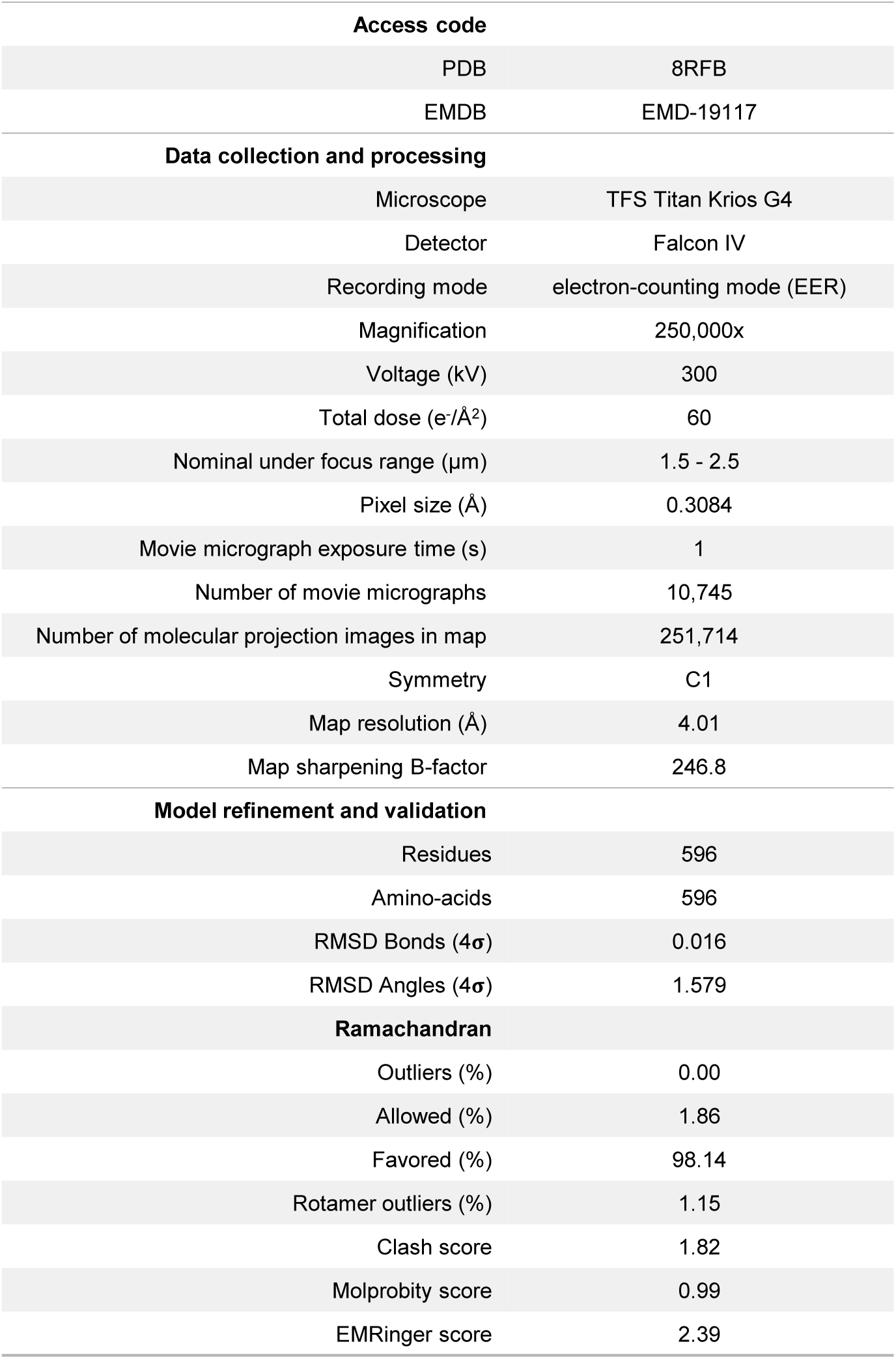
CryoEM map and atomic model refinement.

**Table S2.**
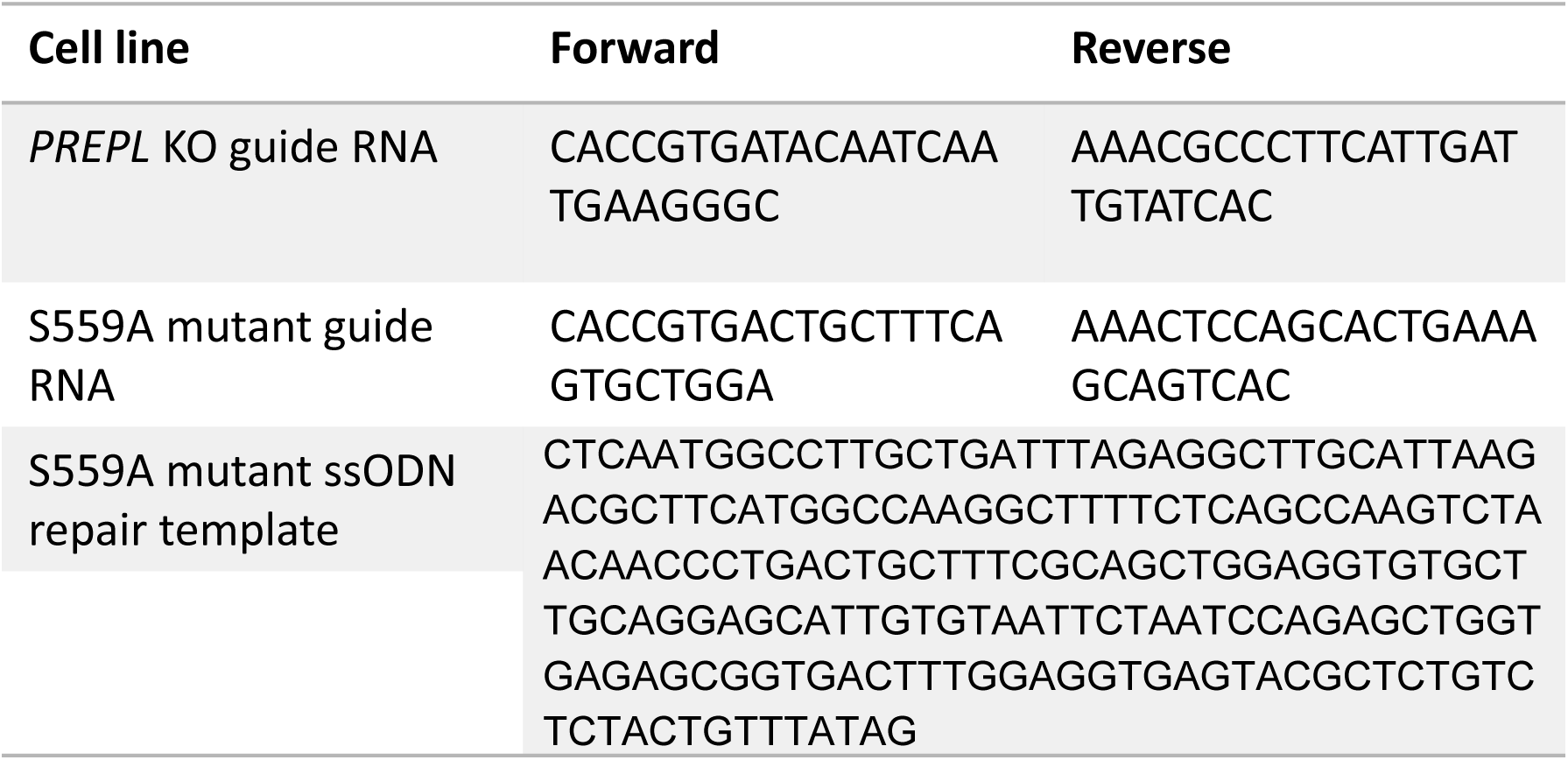
CRISPR-Cas9 oligos.

**Figure S1:**
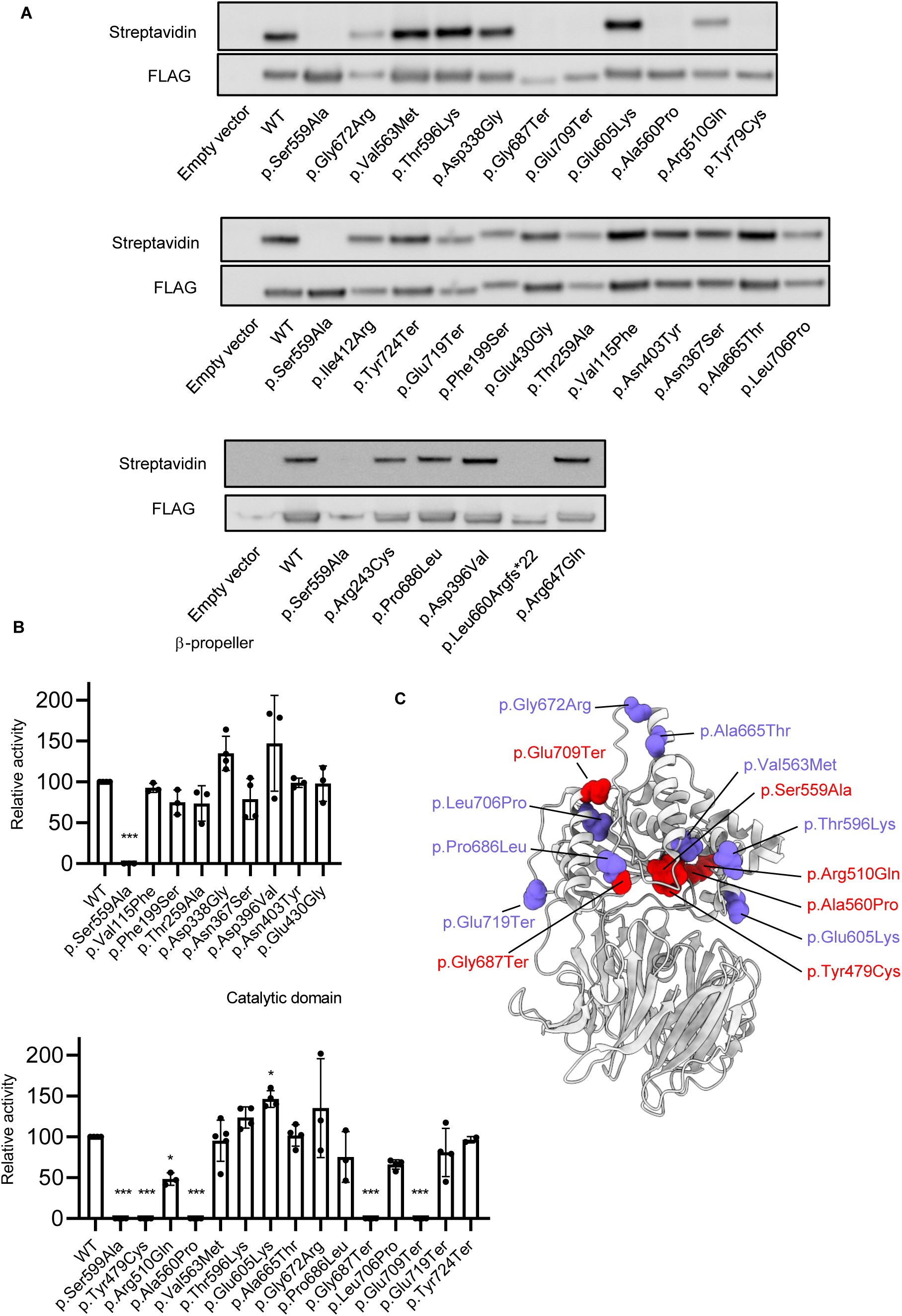
FP-biotin activity-based probe assay on PREPL variants obtained from the ClinVar database. A) FP-biotin – streptavidin blots depicting relative FP-biotin binding. B) Normalised quantitative results of FP-biotin binding. C) Relative location within the protein strucutre of the variants in the catalytic domain. Statistical analysis was performed using one-way ANOVA. Significance levels are shown as * p ≤ 0.05 ** p ≤ 0.01, *** p ≤ 0.001 and **** p ≤ 0.0001

**Figure S2:**
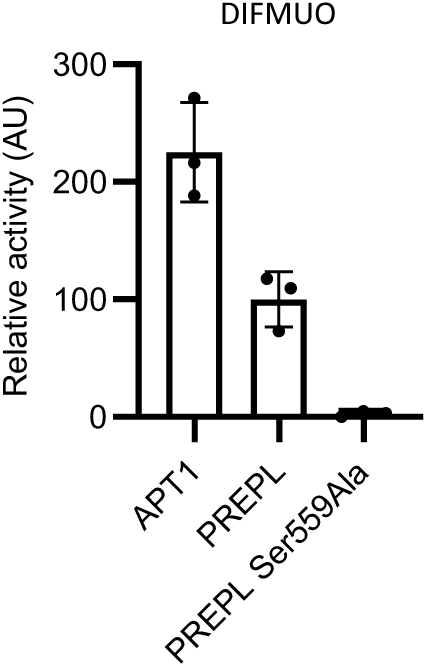
PREPL can cleave DIFMUO at a 50% efficiency compared to APT1. DIFMUO substrate cleavage assay was performed for APT1, PREPL and PREPL p.Ser559Ala variant to validate substrate cleavage capacity of PREPL.

**Figure S3:**
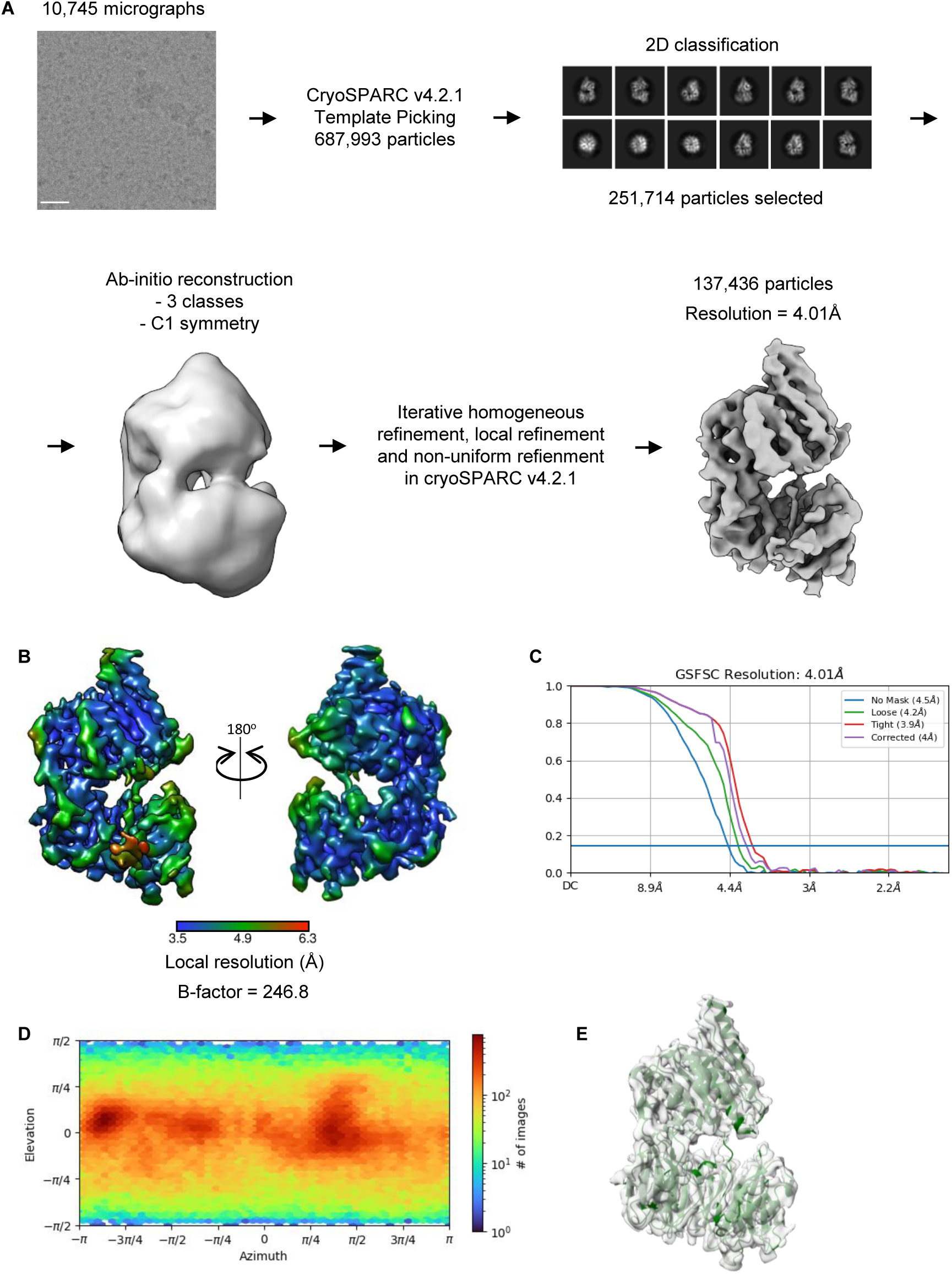
Cryo-EM analysis of PREPL p.Arg243Cys. **(A)** Flow chart of data processing. **(B)** Final 3D reconstruction of PREPL Arg243Cys, coloured according to the local resolution. **(C)** Corrected Gold-standard Fourier shell correlation curves for the 3D electron microscopy reconstruction. **(D)** Angular distribution of PREPL Arg243Cys particles included in the final reconstruction. **(E)** PREPL Arg243Cys model fitted into the map.

**Figure S4:**
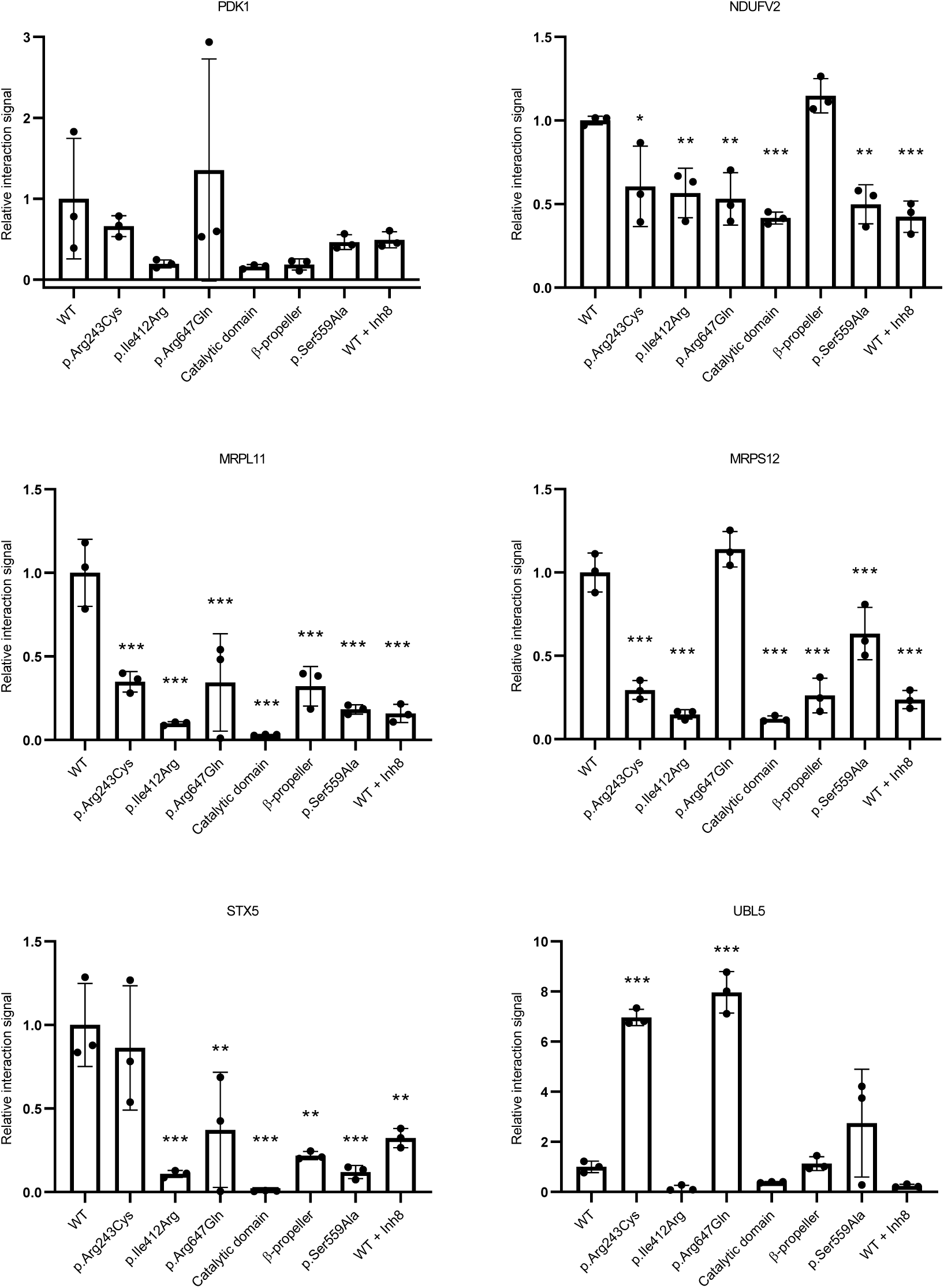
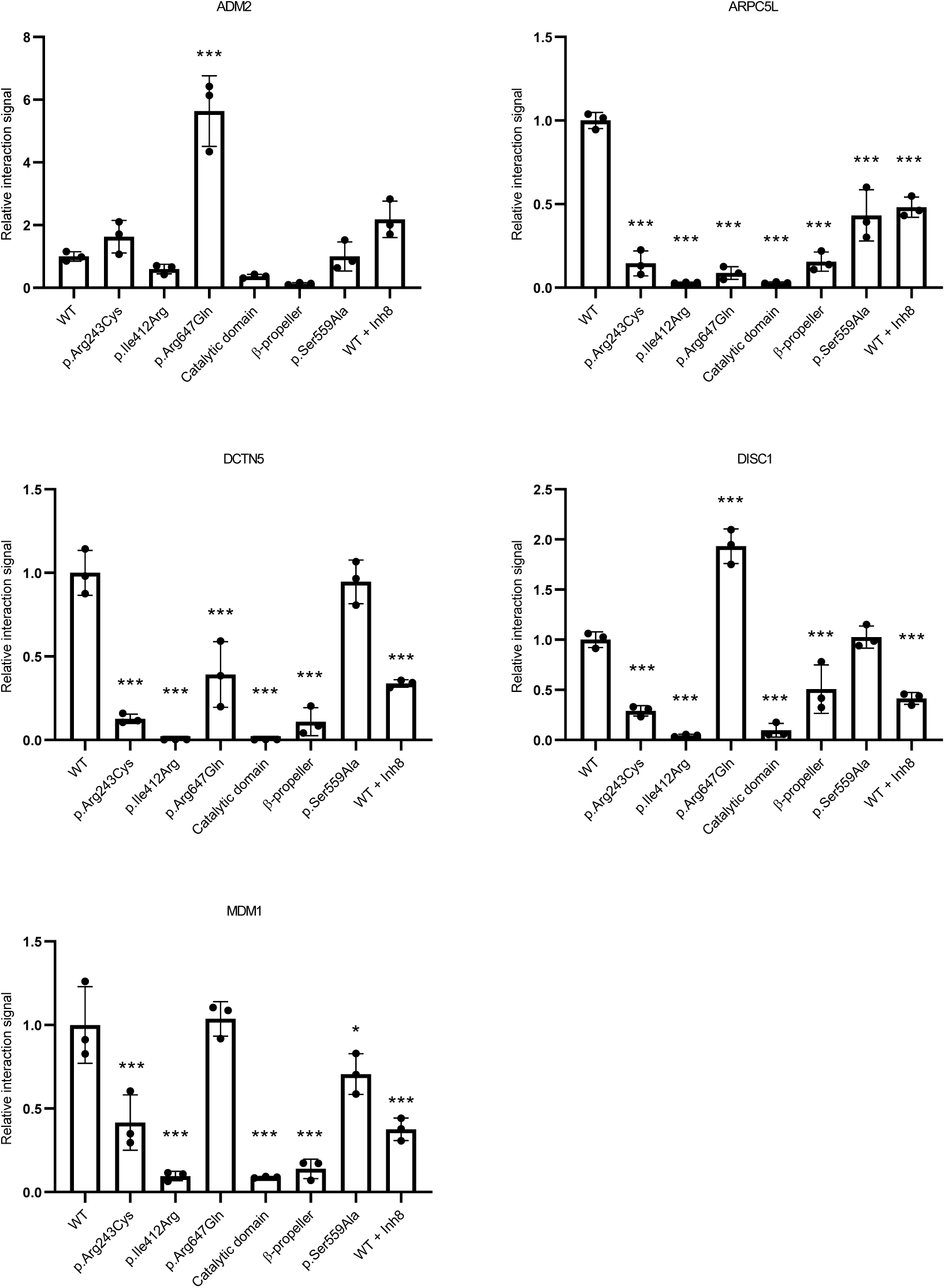
Detailed results of MAPPIT anaylsis of protein-protein interaction scores. Results depicted for each interactor partner with 8 PREPL variants. Statistical analysis was performed using one-way ANOVA. Significance levels are shown as * p ≤ 0.05 ** p ≤ 0.01, *** p ≤ 0.001 and **** p ≤ 0.0001.

**Figure S5:**
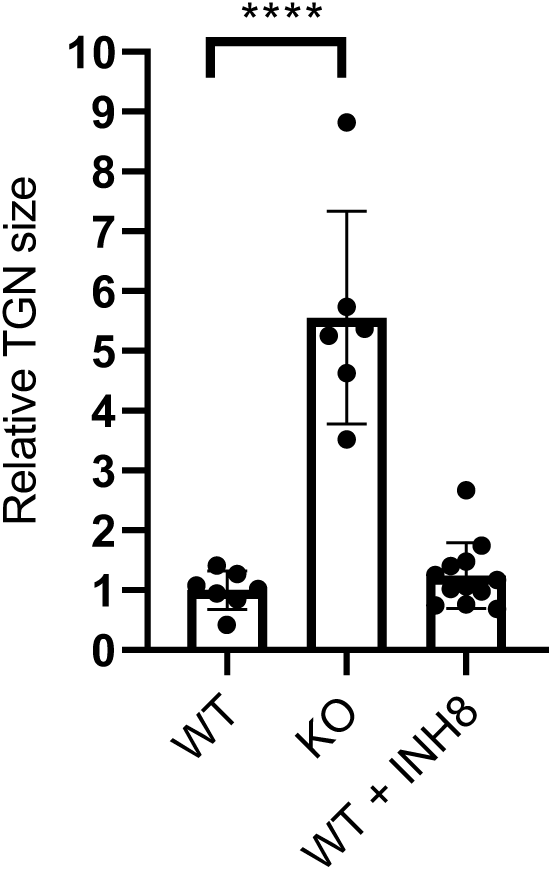
Evaluation of Trans-Golgi network size. Addition of cel-permeable PREPL inhibitor 8 on WT HEK293T cells does not change the relative TNG size compared to non-treated WT HEK293T cells. Statistical analysis was performed by ANOVA. Significance levels are shown as * p ≤ 0.05 ** p ≤ 0.01, *** p ≤ 0.001 and **** p ≤ 0.0001

**Figure S6:**
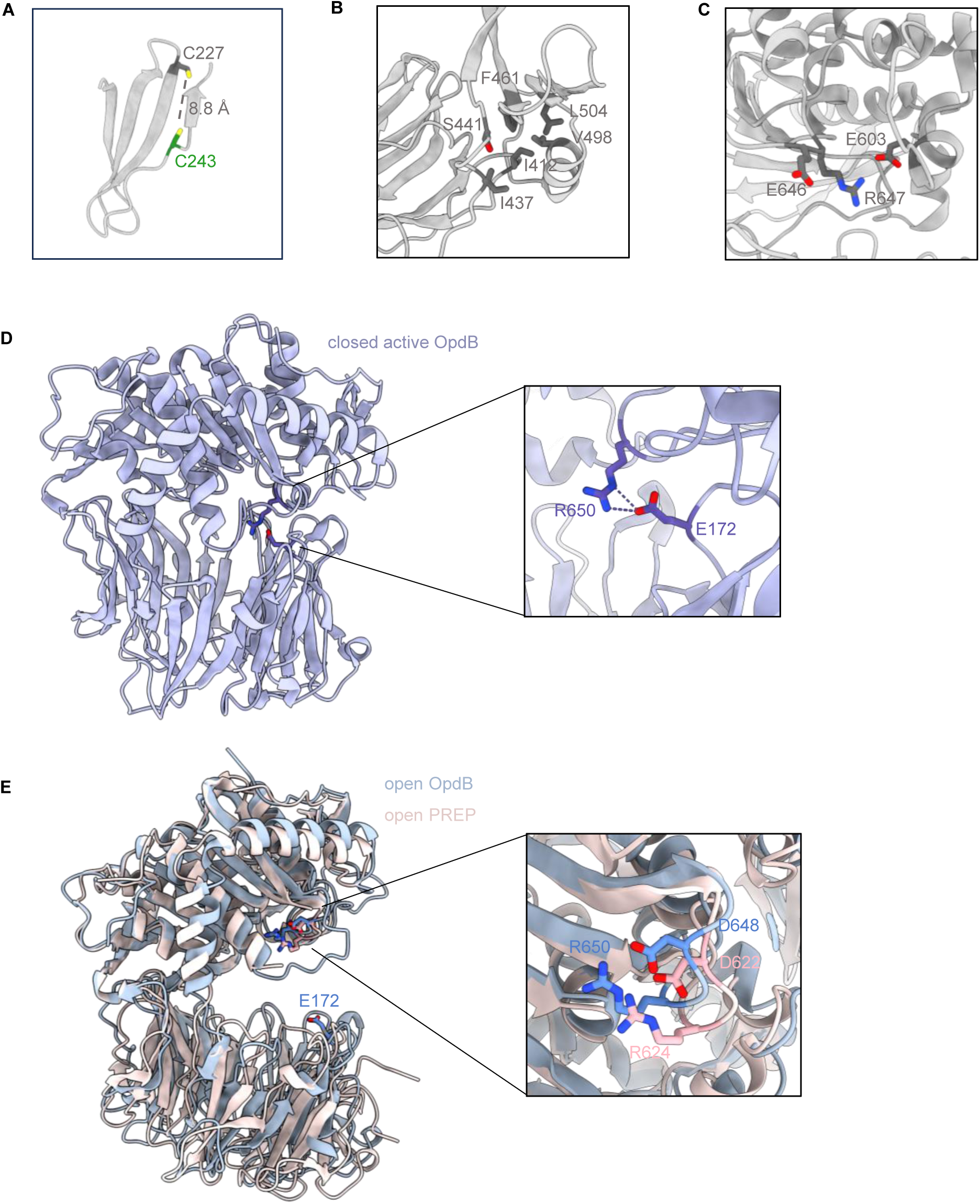
The effect of CMS22 mutations on protein structure. **(A)** The CMS22 Cys243 PREPL mutation and the WT Cys227 on the third blade of the b propeller, found within 9 Å distance. **(B)** The hydrophobic cluster formed on PREPL near Ile412 by Val498, Leu504, Ser441, Ile437, and Phe461. **(C)** Arg647 located near the catalytic domain residues Glu603 and Glu646. **(D)** The closed, active conformation of OpdB (PDB 4BP9), stabilized by the salt bridge formed between the catalytic domain Arg650 (Arg647 in PREPL) and the b-propeller residue Glu172 (Glu180 in PREPL based on structural superposition). **(E)** Superposition of the open conformations of OpdB and PREP, highlighting the interactions formed by Arg650 in OpdB and Arg624 in PREP, the equivalent to PREPL Arg647 residues.

## Notes

https://www.ebi.ac.uk/emdb/

### Competing Interest Statement

The authors have declared no competing interest.

